# Cyclin-dependent-like kinase 5 is required for pain signalling in both human neurons and mouse models

**DOI:** 10.1101/690172

**Authors:** Paolo La Montanara, Arnau Hervera, Lucas Baltussen, Thomas Hutson, Ilaria Palmisano, Francesco De Virgiliis, Yunan Gao, Qasim A. Majid, Nikos Gorgoraptis, Kingsley Wong, Jenny Downs, Vincenzo Di Lazzaro, Tommaso Pizzorusso, Sila Ultanir, Helen Leonard, Nagy Istvan, Nicholas D Mazarakis, Simone Di Giovanni

## Abstract

Cyclin-dependent-like kinase 5 (*Cdkl5)* gene mutations lead to an X-linked disorder that is characterized by infantile epileptic encephalopathy, developmental delay and hypotonia. However, we found that a substantial percentage of these patients also report a previously unrecognised anamnestic deficiency in pain perception. Consistent with a role in nociception, we discovered that Cdkl5 is expressed selectively in nociceptive dorsal root ganglia (DRG) neurons in mice and in iPS-derived human nociceptors. CDKL5 deficient mice display defective epidermal innervation and conditional deletion of *Cdkl5* in DRG sensory neurons significantly impairs nociception, phenocopying CDKL5 deficiency disorder in patients. Mechanistically, Cdkl5 interacts with CaMKIIα to control outgrowth as well as TRPV1-dependent signalling, which are disrupted in both *Cdkl5* mutant murine DRG and human iPS-derived nociceptors. Together, these findings unveil a previously unrecognized role for Cdkl5 in nociception, proposing an original regulatory mechanism for pain perception with implications for future therapeutics in CDKL5 deficiency disorder.

**One Sentence Summary:** Cyclin-dependent-like kinase 5 (Cdkl5) controls nociception in patients and murine models of Cdkl5 deficiency disorder via CaMKII-dependent mechanisms

## Introduction

Mutations in the X-linked *Cdkl5* gene, encoding the cyclin-dependent-like kinase 5, are associated with the CDKL5 Deficiency Disorder (CDD) that is characterized by early-onset intractable seizures, severe intellectual disability, and motor impairment(*1, 2*). Cdkl5 is strongly expressed in the central nervous system (CNS), particularly in neurons of the cortex and of the hippocampus (*3*). Defective Cdkl5 impairs proper brain development, learning and memory, neuronal activity, including molecular processes related to neuronal depolarization and synapse formation (*4–7*). Cdkl5 is activated by neuronal activity and it has also been implicated both in the phosphorylation and interaction with nuclear as well as cytoplasmic proteins potentially modulating gene expression and cytoskeleton changes ultimately affecting neuronal activity and cell survival (*2, 5, 8, 9*). Recently microtubule and centrosome-associated proteins have been identified as targets of Cdkl5 kinase activity, including MAP1S, EB2, ARHGEF2, CEP131 and DLG5, whose regulation is likely to affect axonal transport, outgrowth and synaptic plasticity (*10, 11*).

CDD remains a disease without a cure, whose fundamental molecular and cellular mechanisms are still in need of much investigation and whose full clinical spectrum remains only partially characterized. An important current limitation is that unlike patients, CDD mutant mice do not develop epilepsy(*4, 12–14*). This makes the identification of Cdkl5-dependent mechanisms or potential treatments with an impact on the human phenotype very challenging. Here, we discovered that CDD patients and animal models display an impairment in nociception and that Cdkl5 is required for peripheral nociceptive signalling in dorsal root ganglia (DRG) neurons in mice and in induced pluripotent cell (iPS)-derived human nociceptors. Mechanistically, conditional deletion of Cdkl5 in DRG sensory neurons significantly impairs nociception, phenocopying CDKL5 deficiency disorder in patients. Lastly, we discovered that Cdkl5 interacts with CaMKIIα to control outgrowth of sensory neurons as well as TRPV1-dependent signalling including in iPS-derived neurons from CDD patients. Together, these findings reveal a previously unrecognized role for Cdkl5 in nociception and they suggest an original peripheral regulatory mechanism for pain processing with implications for future therapeutic interventions in CDD.

## Results

### Alteration in nociception in CDD patients

In a recent effort to broadly characterise the clinical phenotype, variation and natural history of CDD, the International CDKL5 Database (ICDD) (https://www.cdkl5.com/cdkl5-international-registry-database/) was established. Analysis of this clinical data source allowed the identification of a previously unrecognised significant occurrence of alterations in pain perception. In fact, the database included responses to questions to caregivers about whether their child currently experiences alterations in pain perception.

Since nociception had not been characterized so far in CDD and it can be promptly clinically assessed in both humans and animal models, we decided that it deserved further investigation. Patients were classified by age group and mutation type (Table 1). Classification of individual CDKL5 mutations was based on predicted structural and functional phenotypic consequences similar to the groupings used in a previous study (*15*). In our analysis we have grouped together mutations leading to lack of functional protein and missense/in-frame mutations within catalytic domain and TEYmotif and compared them with mutations affecting the regulatory domains of *Cdkl5* i.e. truncations after aa172. Binary regression with log link was used to estimate the relative risk of altered pain sensitivity including by age groups and mutation type (Table 1).

We found that altered pain sensitivity in their children was reported by 53.0% (122/230) of caregivers. Amongst these a total of 57.4% (70/122) specifically reported decreased pain sensitivity, 19.7% (24/122) specifically reported enhanced sensitivity and 22.9% (28/122) reported both. 202 caregivers provided specific responses including either reduced (70/202, 34.7%) or enhanced nociception (24/202, 11.9%) (Table 1). Amongst those who reported either reduced or enhanced pain sensitivity, a total of 74.4% (70/94) reported decreased and 25.5% (24/94) increased nociception. The likelihood of the response being specifically reduced pain perception was measured as risk ratio (RR) of reduced or enhanced pain sensitivity versus normal pain perception. As shown in Table 1 reduced pain perception was more likely to be reported than enhanced pain perception (RR 2.92, 95% CI 2.89,7.00; p<0.001). Compared with those aged two years or under, individuals aged six years and over were reported to have an increased likelihood of reduced sensitivity to pain (RR 2.74, 95% CI 1.36,5.53; p=0.005). Moreover, compared with those with mutations in the *Cdkl5* regulatory domain (truncations after aa172), those with a non-functional protein (including missense/in-frame mutations within catalytic domain and TEY motif) were also reported to have an increased likelihood of reduced sensitivity to pain (RR 1.63, 95% CI 0.87,3.06; p=0.124). Reduced was significantly more likely than enhanced pain perception at each age group (0-2 years old, (RR 3.00, 95% CI 1.28,7.06; p=0.012); 3-5 years old, (RR 2.67, 95% CI 1.04,6.81; p=0.040); 6+ years old, (RR 3.00, 95% CI 1.52,5.94; p=0.002) and for each gene mutation group studied (no functional protein, (RR 2.94, 95% CI 1.67,5.18; p<0.001); truncation after aa172, (RR 2.88, 95% CI 1.29,6.42; p=0.010).

**Table 1.** Associations between pain sensitivity and age group and mutation type in 202 individuals with confirmed CDKL5 mutation.

|  |  | None | Reduced | Reduced versus None |  | Enhanced | Enhanced versus None |  | Reduced versus Enhanced |
| --- | --- | --- | --- | --- | --- | --- | --- | --- | --- |
|  |  |  |  | Within category | Between category |  | Within category | Between category | Within category |
|  | N | N (%) | N (%) | RR (95% CI) | RRR (95% CI) | n (%) | RR (95% CI) | RRR (95% CI) | RR (95% CI) |
| **Total** | 202 | 108 (53.5) | 70 (34.7) | 0.65 (0.48,0.88) | n/a | 24 (11.9) | 0.22 (0.14,0.35) | n/a | 2.92 (2.89,7.00) |
| **Age group (years)** |  |  |  |  |  |  |  |  |  |
| 0-2 | 82 | 54 (65.9) | 21 (25.6) | 0.39 (0.23,0.64) | 1.00 | 7 (8.5) | 0.13 (0.06,0.28) | 1.00 | 3 (1.28,7.06) |
| 3-5 | 45 | 23 (51.1) | 16 (35.6) | 0.70 (0.37,1.32) | 1.79 (0.79,4.03) | 6 (13.3) | 0.26 (0.11,0.64) | 2.01 (0.61,6.65) | 2.67 (1.04,6.81) |
| 6+ | 75 | 31 (41.3) | 33 (44.0) | 1.06 (0.65,1.73) | 2.74 (1.36,5.53) | 11 (14.7) | 0.33 (0.17,0.66) | 2.74 (0.96,7.79) | 3 (1.52,5.94) |
| **Mutation type** |  |  |  |  |  |  |  |  |  |
| Truncations after aa172 | 79 | 48 (60.8) | 23 (29.1) | 0.48 (0.29,0.79) | 1.00 | 8 (10.1) | 0.17 (0.08,0.35) | 1.00 | 2.88 (1.29,6.42) |
| No functional protein* | 123 | 60 (48.8) | 47 (38.2) | 0.78 (0.53,1.15) | 1.63 (0.87,3.06) | 16 (13.0) | 0.27 (0.15,0.46) | 1.6 (0.63,4.05) | 2.94 (1.67,5.18) |

None: patients with normal pain sensitivity; reduced or enhanced: individuals with reduced or enhanced pain sensitivity
N, number of individuals; RR, relative risk; RRR, relative risk ratio; CI, confidence interval; Ref, reference category
Note: excluded individuals where both reduced and enhanced sensitivity were reported
\* including missense/in-frame mutations within catalytic domain and TEY motif

### Cdkl5 localizes to human and murine sensory neurons and is required for nociception

Since primary nociceptors are localized in the dorsal root ganglia, we initially investigated whether Cdkl5, so far localised to the CNS, was also expressed in the peripheral nervous system in murine as well as human nociceptors. Surprisingly, immuno and co-immunohistochemistry studies in DRG revealed that Cdkl5 is indeed expressed and mainly found in the cytoplasm of small diameter DRG neurons (Fig. 1A-C), where the signal becomes detectable after the age of post-natal day P45 (fig. S1A,B). Around 90% of Cdkl5 positive neurons express the nociceptive markers CGRP or IB4 (Fig. 1D-F). The expression of *Cdkl5* in murine DRG was confirmed by RT-PCR and immunoblotting (fig. S1C,D), where *Cdkl5* was expressed although to a lower level compared to the brain.

**Figure 1.**
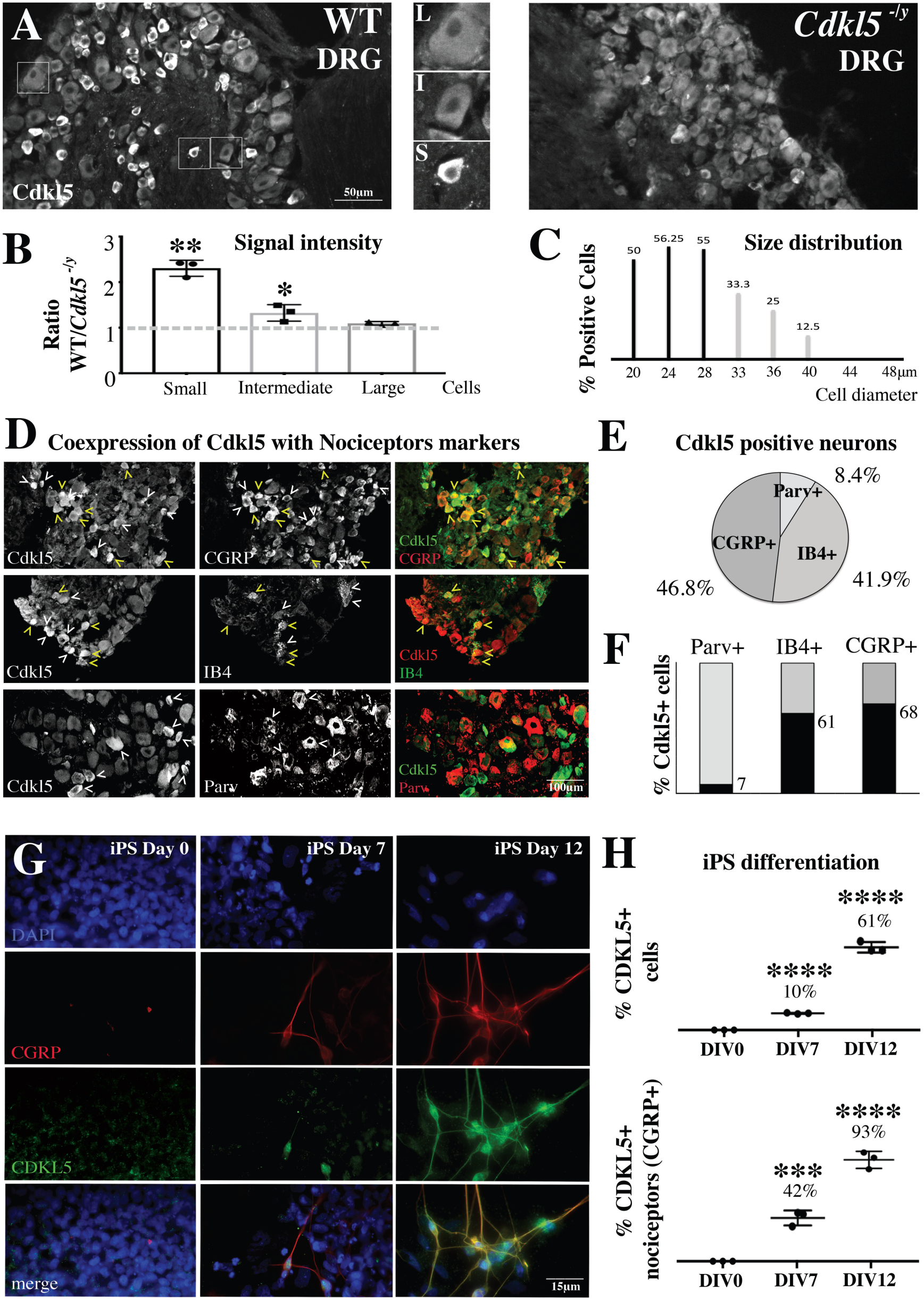
Cdkl5 is expressed in nociceptors. **A.** Cdkl5-immunostaining in DRG from WT and Cdkl5^-/y^ mice (20x, Nikon Eclipse-TE-2000U microscope). L-I-S: insets show high magnification of large, intermediate and small diameter DRG neurons. **B**. Graphs showing ratio of signal intensity between WT and *Cdkl5^-/y^* three neuronal subpopulations. N: 3 mice each, Mean with SD, Student’s *t-*test, \*\**P<0.01 *P<0.05.* **C**. Graphs showing percentage of Cdkl5+ positive neurons in the indicated sizes. N:3 mice each. **D**. Confocal immunofluorescence images of DRG biopsies from P70 WT mice showing the co-expression of Cdkl5, CGRP, IB4 and Parvalbumin (20x images from Zeiss LSM-780; yellow arrowheads: co-expression; white arrow heads: expression of Cdkl5 or individual cell type markers only). **E**. Percentage of Cdkl5+ cells co-expressing each marker (average percentage values represent average from 3 independent mice). **F**. Percentage of Parvalbumin+, IB4+ or CGRP+ cells co-expressing Cdkl5+ (average percentage values represent average from 3 independent mice). **G**. CDKL5 and CGRP immunostaining from isogenic iPS derived neurons after 0, 7 and 12 days of differentiation into nociceptors (63x, Leica TCS SP5II). **H**. Graphs showing the percentage of CDKL5+ cells and CDKL5+ neurons coexpressing CGRP at different stages of differentiation. N: 3, Mean with SD, One way ANOVA, Tuckey’s post-hoc, **\*\*\*\****P<0.0001* \*\*\**P<0.001*.

Next, in order to find whether Cdkl5 was also expressed in human nociceptors, we differentiated iPSC obtained from human skin biopsies into nociceptors. After having confirmed the differentiation of iPSC by immunolabelling for βIII-Tubulin as neuronal and CGRP as nociceptor marker respectively (fig. S2A-E), we observed that Cdkl5 was indeed expressed in human nociceptors (Fig. 1G,H).

In order to directly address whether the expression of *Cdkl5* in DRG sensory neurons was required for nociception, we took advantage of the conditional deletion of Cdkl5 in sciatic DRG neurons by injecting an AAV5-creGFP or control GFP virus to the sciatic nerve of *Cdkl5*-floxed mice (*12*) (Fig. 2A). Since the cre-virus deletes Cdkl5 selectively in DRG neurons, the absence of Cdkl5 signal by immunoblotting strongly supports the selective neuronal expression in DRG ganglia as well as the efficiency of the deletion (Fig. 2B). Cdkl5 conditionally deleted mice showed an increase in paw withdrawal latency and threshold in response to noxious hot temperature (*Haargraves plantar test)* and noxious mechanical stimulation *(Von Frey)* respectively (Fig. 2C,E). We performed the same functional tests in *Cdkl5* mutant mice, an established mouse model of CDD carrying a nonsense mutation (stop codon) leading to deletion of *Cdkl5*-Ex6 (*4*). Here, we found a reduction in nociception compared to WT littermates that phenocopied what observed in *Cdkl5* conditionally deleted mice (Fig. 2D,F). Importantly, in Cdkl5 conditionally deleted mice additional sensory modalities were unaffected as indicated by the time-to-contact and the time-to-removal during the adhesive removal test (*16*) (Fig. 2G). Sensorimotor functions as assessed by the gridwalk task (*17*), which evaluates motor coordination and proprioception were also unaltered (Fig. 2H), suggesting a selective role for CDKL5 in nociception. Together, these data suggest that Cdkl5 is required for physiological nociception that relies upon the expression of Cdkl5 in DRG neurons.

**Figure 2.**
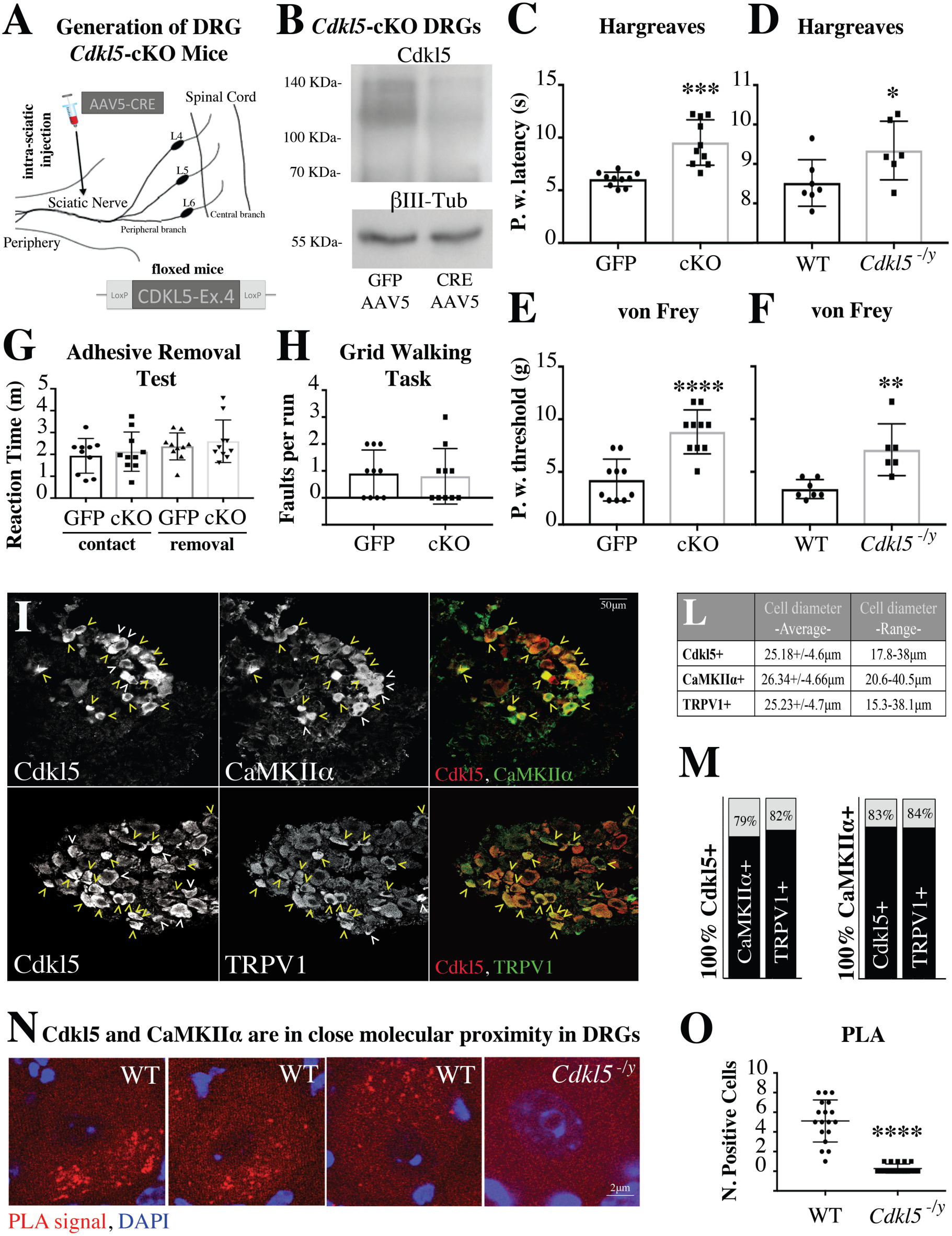
Conditional deletion of *Cdkl5* impairs nociception. **A.** Schematic illustrating conditional deletion of *Cdkl5* in DRG neurons. Depicted is the injection of AAV5 particles into the sciatic nerve of 12 weeks old *Cdkl5*-floxed mice. **B**. Immunoblotting showing strongly reduced expression of Cdkl5 from bilateral sciatic DRGs (pool of 4 mice) 8 weeks after AAV5-GFP or AAV5-GFP-cre. βIII-tubulin has been used as loading control. **C-F**. Behavioral assessment of pain sensitivity with Hargreaves (thermal nociception; Paw withdrawal latency) and von Frey (mechanical allodynia; Paw withdrawal threshold) shows significantly reduced sensitivity in *Cdkl5* conditionally deleted (**C, E**) as well as in *Cdkl5^-/y^* mice (**D, F**). **C, E**. N:10 mice, Mean with SD, Student’s *t*-test, \*\*\*\**P<0.0001* \*\*\**P<0.001*. **D,F**. N: 7 WT vs 6 Mut mice, Mean with SD, Student’s *t*-test, \*\**P<0.01 *P<0.05*. **G**. Graph showing no difference in the reaction time (time to contact and time to remove the adhesive tape from the paw) in Cdkl5 DRG conditionally deleted (cKO) vs control GFP injected mice. N:10, Mean with SD, Student’s t-test, *P>0.05*. **H**. Graph showing no difference in the number of posterior foot faults in each run on a Gridwalk in Cdkl5 DRG conditionally deleted (cKO) vs control GFP injected mice. N: 10 mice per group, Mean with SD, Student’s t-test, *P>0.05*. **I**. Confocal microscopy of immunofluorescence showing co-expression of Cdkl5, CaMKIIα and TRPV1 (magnification 20x, Leica TCS SP5II; yellow arrowheads: co-expression; white arrowheads: expression of Cdkl5 or individual cell type markers only). **L**. Average diameter and SD of each Cdkl5+, CaMKIIα+ or TRPV1+ population of cells (average values from 3 independent mice). **M**. Percentage of Cdkl5+ or CaMKIIα+ cells co-expressing each marker (average percentage values represent average from 3 independent mice. Cdkl5+ cells coexpressing CaMKIIα: 79.07+/- 1.56%; Cdkl5+ cells coexpressing TRPV1: 82.3+/-6.8%; CaMKIIα+ cells coexpressing Cdkl5: 83.97+/-3.44%; CaMKIIα+ cells coexpressing TRPV1: 84.01+/-5.66%. **N**. Confocal images showing individual WT or *Cdkl5^-/y^* DRG neurons (63x, Leica TCS SP5II) after proximity ligation assay (PLA) (red) and DAPI staining (blue). The presence and intensity of red dots represent the presence and degree of molecular proximity between Cdkl5 and CaMKIIα. **O**. Graph showing the number of PLA-positive cells in WT and *Cdkl5^-/y^* DRGs. N: average number of positive DRG cells from 3 WT and 3 *Cdkl5^-/y^* mice, Mean with SD, Student’s *t*-test, \*\*\*\**P<0.0001*.

### Cdkl5 controls the outgrowth of human and murine sensory neurons via a CaMKII-dependent mechanism

To investigate the signalling pathways involved in Cdkl5-dependent pain transmission, we decided to identify interactors of Cdkl5 *in vivo*. To this end, we performed immunoprecipitation of Cdkl5 from the murine cortex, where the kinase is highly enriched, followed by mass spectrometry of the eluate (Suppl. File 1 and fig. S3A). Considering the normalized protein ratios of each immunoprecipitation experiment (n=6), we identified 23 proteins co-immunoprecipitating with Cdkl5 (fig. S3B). Interestingly, 74% of them correspond to genes associated with neuronal activity, neural development and epilepsy (fig. S3C-F), supporting the physiological relevance of our pool of interactors. Top ranked Cdkl5 co-immunoprecipitating proteins included CaMKIIα, putatively the strongest interactor of Cdkl5, and some proteins associated with the neuronal cytoskeleton, such as Myh10 (*18*), Tubb3 (*19*) and Dynch1h1 (*20–23*) (fig. S3G). The association with CaMKIIα suggests also a role for Cdkl5 in calcium-dependent signalling, neuronal activity (*24*) and in cytoskeleton remodelling (*25–30*).

Previous immunohistochemical studies have found that CaMKII is expressed in DRG nociceptors (*31*), where it plays a role in the regulation of neurite outgrowth (*32*). In particular, CaMKII is expressed in TRPV1-immunoreactive nociceptors, in both CGRP+ and IB4+ subpopulations, where it is required for capsaicin-mediated nociception via modulation of TRPV1; capsaicin stimulation of TRPV1 and calcium entry activate CaMKII that is in turn required to potentiate TRPV1 signalling (*33, 34*). Hence, we hypothesised that Cdkl5 could be a partner of CaMKIIα in the regulation of pain both via modulation of cytoskeleton remodelling and nociception signalling pathways.

First, we examined by confocal microscopy the co-expression of Cdkl5, CaMKIIα and TRPV1 in DRG neurons and we found that they co-localize in most DRG neurons (Fig. 2I-M). Next, we confirmed, by co-immunoprecipitation and immunoblotting from brain extracts, that Cdkl5 does interact with CaMKIIα in vivo (fig. S3H). Similarly, co-immunoprecipitation experiments after overexpression of CDKL5 in HEK293 cells, or of its inactive kinase dead form CDKL5-K42R and CaMKIIα, confirmed the interaction between the active form of CDKL5 and CaMKIIα. Importantly, no interaction was observed between CaMKIIα and CDKL5-K42R (fig. S3I), which showed lack of kinase activity in an in vitro kinase assay (fig. S3J), supporting the specificity of the findings. We finally established the molecular proximity between Cdkl5 and CaMKIIα in DRG neurons by using the Proximity Ligation Assay (Fig. 2N,O), supporting the co-expression and co-immunoprecipitation data.

We next investigated whether Cdkl5-CaMKIIα signalling axis would be required for the outgrowth of DRG neuronal processes. Assessment of neurite outgrowth was carried out in both murine and human iPSC derived nociceptors from *Cdkl5* null mice and CDKL5 deficiency disorder patients. We found that cultured DRG neurons from *Cdkl5* mutant mice display impaired neurite outgrowth in CGRP positive neurons (Fig. 3A,C), and this is rescued to the WT levels by viral-mediated overexpression of CDKL5 (Fig. 3E; fig. S4). On the contrary, outgrowth in proprioceptive parvalbumin positive neurons remained unaffected by Cdkl5 loss of function (Fig. 3B,D), indicating a specific role for Cdkl5 in nociceptors. Importantly, impaired neurite outgrowth was also observed in iPSC-derived nociceptive DRG neurons (CGRP+) obtained from skin biopsies of CDKL5 patients, compared with the respective isogenic controls (Fig. 3F,G). CGRP+ neurons represent 30-40% of WT and mutant DRG neurons and more than 90% of iPSC derived sensory neurons were βIII-Tubulin+ (fig. S2D). Capsaicin responding cells were ∼60% of the KCl responding murine DRG cultured neurons, while they were more than 90% of the KCl responding iPSC derived neurons, both in mutant and isogenic cultures, indicating functionality of the nociceptors (fig, S2E). Interestingly, we found that neurite outgrowth was impaired in neurons derived from patients carrying a *Cdkl5* mutation compromising the kinase active site and the TEY motif, but not in neurons where only the terminal part of the Cdkl5 tail is lost (Fig. 3H; Table 1).

**Figure 3.**
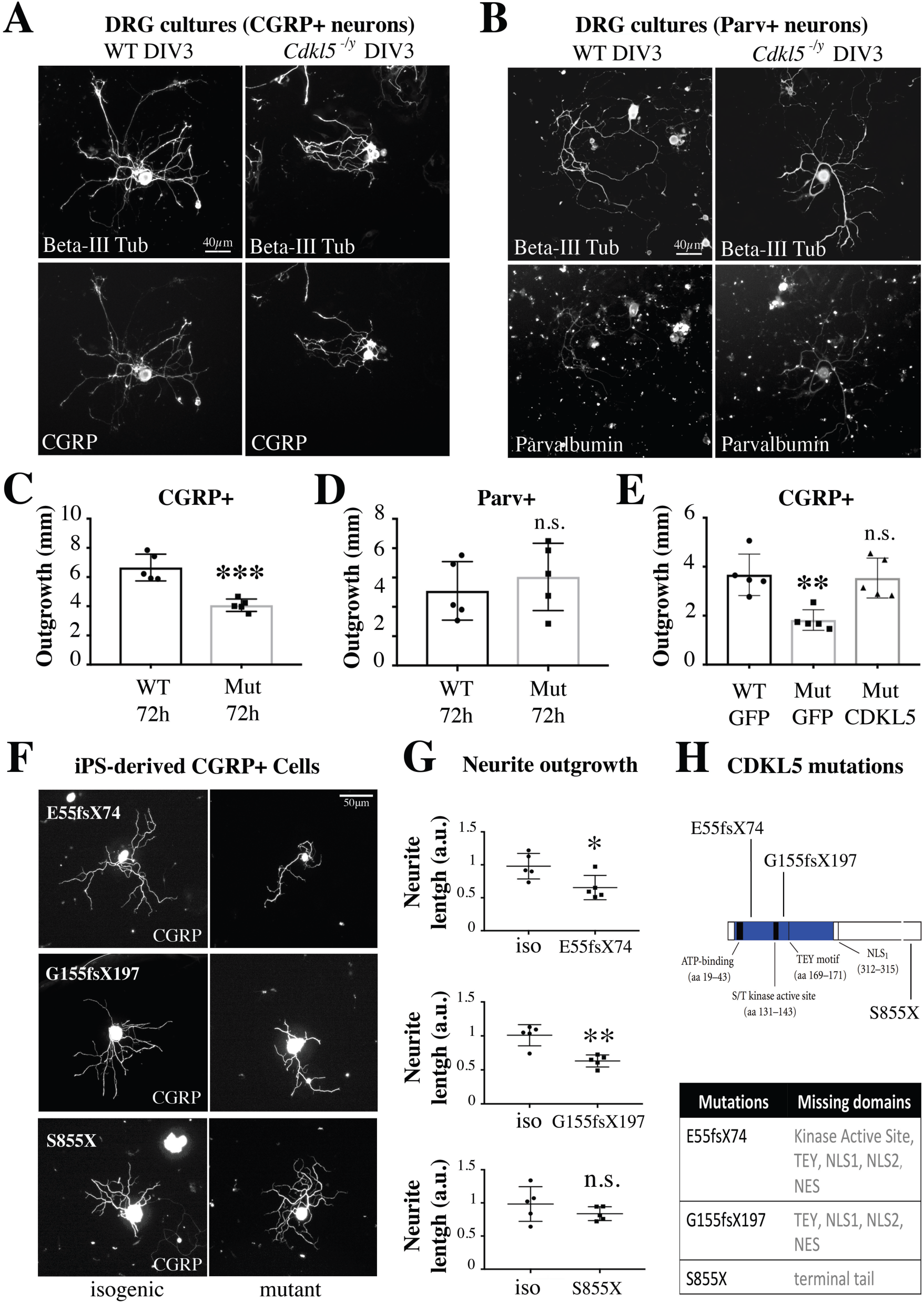
Cdkl5 is required for outgrowth of human sensory neurons. **A,B.** Representative immunofluorescence of βIII-tubulin, CGRP or Parvalbumin in WT and *Cdkl5^-/y^* DRG cultured neurons (DIV3) showing reduced neurite outgrowth in CGRP+ but nor Parv+ *Cdkl5^-/y^* neurons. **C.** Graph showing reduced neurite outgrowth in CGRP+ neurons from *Cdkl5^-/y^* mice (N:5 biological replicates, Mean with SD, Student’s *t*-test, \*\*\**P<0.001*) that is rescued by overexpression of CDKL5 (**E**; N: 5 biological replicates, Mean with SD, One way ANOVA, Tukey’s post-hoc, \*\**P<0.005*). **D.** Graph showing equal neurite outgrowth in Parv+ neurons from WT and *Cdkl5^-/y^* mice. N: 5 biological replicates, Mean with SD, Student’s *t*-test, *P>0.05*. **F.** Representative images of CGRP immunostaining of iPS derived sensory neurons from three CDKL5 patients carrying unique mutations (E55fsX74; G155fsX197; S855X), compared to their isogenic controls (20x images, Nikon TE-2000U fluorescence microscope). **G**. CGRP immunostaining was used to measure total average neurite outgrowth (NeuronJ) of nociceptive neurons. N: 5 independent experiments, total neurite outgrowth/cell. Mean with SD, Student’s *t-* test, **\****P<0.05 **P<0.01*. **H**. Schematic representation of human CDKL5 protein (adapted, (*2*) with the catalytic domain in blue (ATP-binding site, aa 19-43; kinase active site, aa 131-143; TEY motif, aa 169-171) and the C-terminal tail in white (Nuclear Localization Signal 1, aa 312-315; Nuclear Localization Signal 2, aa 784-789; Nuclear Export Signal, aa 836-845).

Next, we asked whether CaMKII activity was required for Cdkl5-dependent neurite outgrowth. Indeed, we found that DRG outgrowth was significantly impaired following administration of the specific CaMKII enzymatic inhibitor KN93, while its inert structural analog KN92 did not have an effect (Fig. 4A,B). Interestingly, KN93 did not further reduce outgrowth in *Cdkl5* mutant cells (Fig. 4A,B), indicating that CDKL5 and CaMKII belong to the same signalling pathway. Importantly, CDKL5 overexpression rescued outgrowth defects in *Cdkl5* mutant knockout DRG neurons, but this effect was blocked by KN93. These data together suggest that Cdkl5-dependent DRG outgrowth relies on CaMKII.

**Figure 4.**
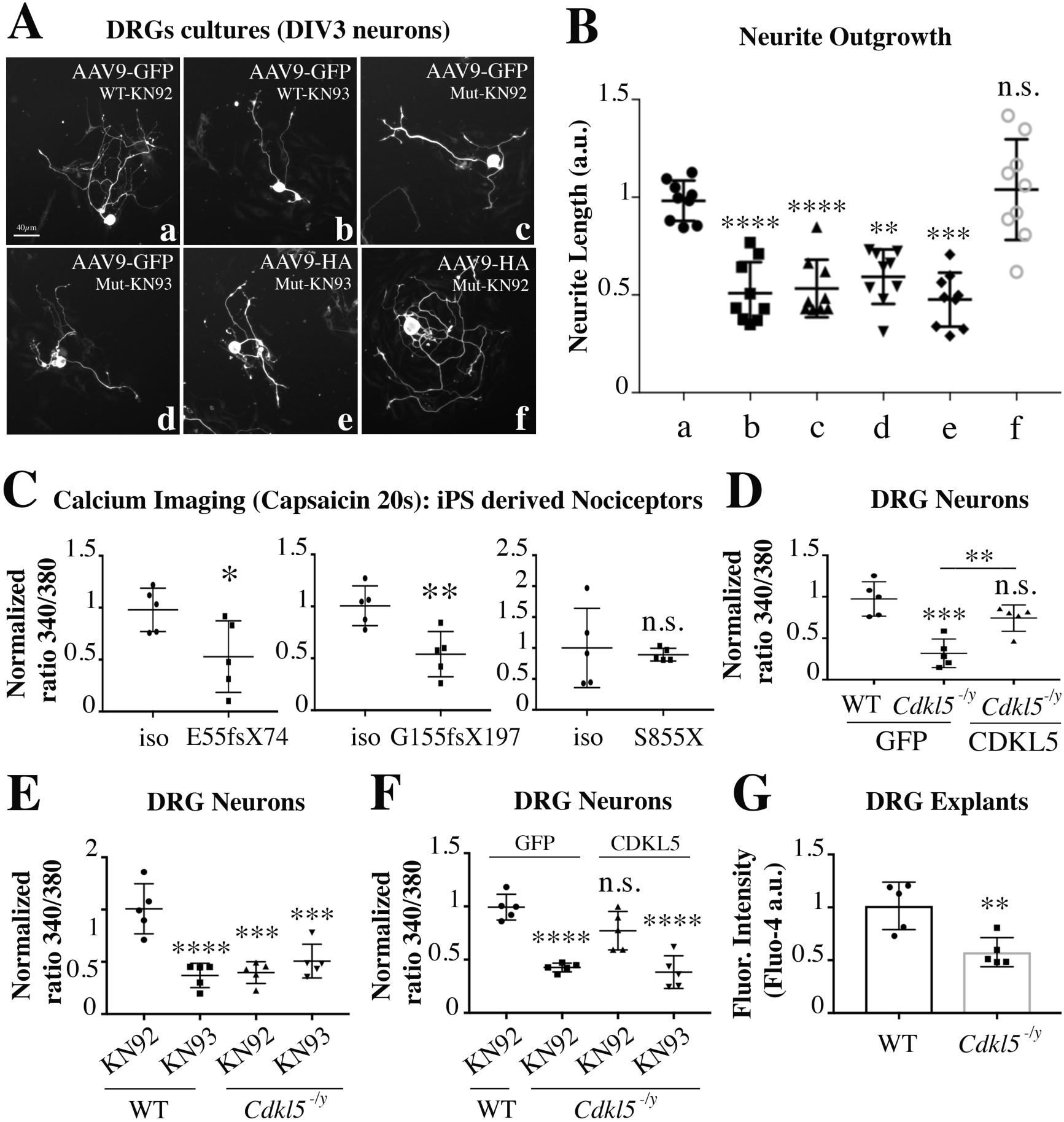
Cdkl5/CaMKII signalling is required for capsaicin signalling and outgrowth of sensory neurons. **A.** Representative immunofluorescence of DRG cultured neurons (DIV3, from P70 WT and *Cdkl5^-/y^* mice), immunostained for GFP or HA after neurons had been infected with AAV9 particles expressing GFP or HA-CDKL5_107_ at DIV1 (20x, Nikon TE-2000U fluorescence microscope). DRG neurons were also treated with 0.5µM of the CaMKII inhibitor KN93 or its inert structural analog KN92. **B**. Graphs showing neurite outgrowth of GFP or HA positive cells that was normalized to the ctrl WT-GFP. N: 9, biological replicates, Mean with SD, One way ANOVA, Tukey’s post-hoc, **\*\*\*\****P<0.0001* \*\*\**P<0.001* \*\**P<0.005*. **C**. Calcium imaging (Fura-2) from iPS derived nociceptors carrying three different CDKL5 mutations (E55fsX74; G155fsX197; S855X), compared with the respective isogenic controls. Graphs showing quantification of intracellular calcium levels in response to capsaicin. E55fsX74 and G155fsX197 neurons show reduced calcium influx after capsaicin. N: 5, average single cell Fura-2 normalised signal/well (∼30 cells per well). Mean with SD, Student’s *t-*test, \*\**P<0.01* \**P<0.05*. **D-F**. Calcium imaging in cultured DRG neurons (Fura-2). DIV3 *Cdkl5^-/y^* neurons showing reduced calcium influx after capsaicin and rescue to WT levels after overexpression of CDKL5 (sspTR-CBh-HA-CDKL5_107_ vs GFP AAV9 particles, infection at DIV1) (**D**). The same experimental conditions as (D) after pre-treatment with KN93 or KN92 in WT and *Cdkl5^-/y^* neurons (**E**), overexpressing CDKL5 or GFP (**F**). N: 5, average single cell Fura-2 normalised signal /well (∼30 cells per well), Mean with SD, One way ANOVA, Tukey’s post-hoc, \*\*\*\**P<0.0001* \*\*\**P<0.001* \*\**P<0.005.* **G**. Calcium imaging in DRG-explants from WT and *Cdkl5^-/y^*mice. Graphs showing quantification of intracellular calcium levels in response to capsaicin (Fluo-4). N: 5, average single cell Fura-4 normalised signal /well (∼30 cells per well), Mean with SD, Student’s *t-*test, \*\**P<0.01*.

Since epidermal innervation is needed for proper nociception and it is a dynamic process requiring continuous cytoskeleton remodelling and outgrowth, we examined if Cdkl5 deficiency was associated with a defect in epidermal innervation.

Indeed, immunohistochemical analysis of skin innervation revealed impairment in epidermal but not dermal innervation in *Cdkl5* mutant versus WT adult mice (Fig. 5A-E). We also measured epidermal innervation from the post-natal day 4 (P4) to adulthood (P70) and found that *Cdkl5* mutant mice display deficient epidermal innervation only in adulthood (P70), and not at earlier stages (P4 or P16; Fig. 5B). In support of these data, we found, by immunoblotting of skin lysates, that the specific neuronal and axonal proteins PGP9.5, CGRP and βIII-tubulin are reduced in *Cdkl5* mutants as compared to WT adult mice (Fig. 5F,G). However, consistent with the selective reduction of nociceptive fibres in the epidermis of *Cdkl5* mutant mice, the expression of these axonal proteins is not reduced in sciatic nerve lysates (Fig. 5H,I) or after immunostaining for CGRP (Fig. 5J,K), in *Cdkl5* mutant mice.

**Figure 5.**
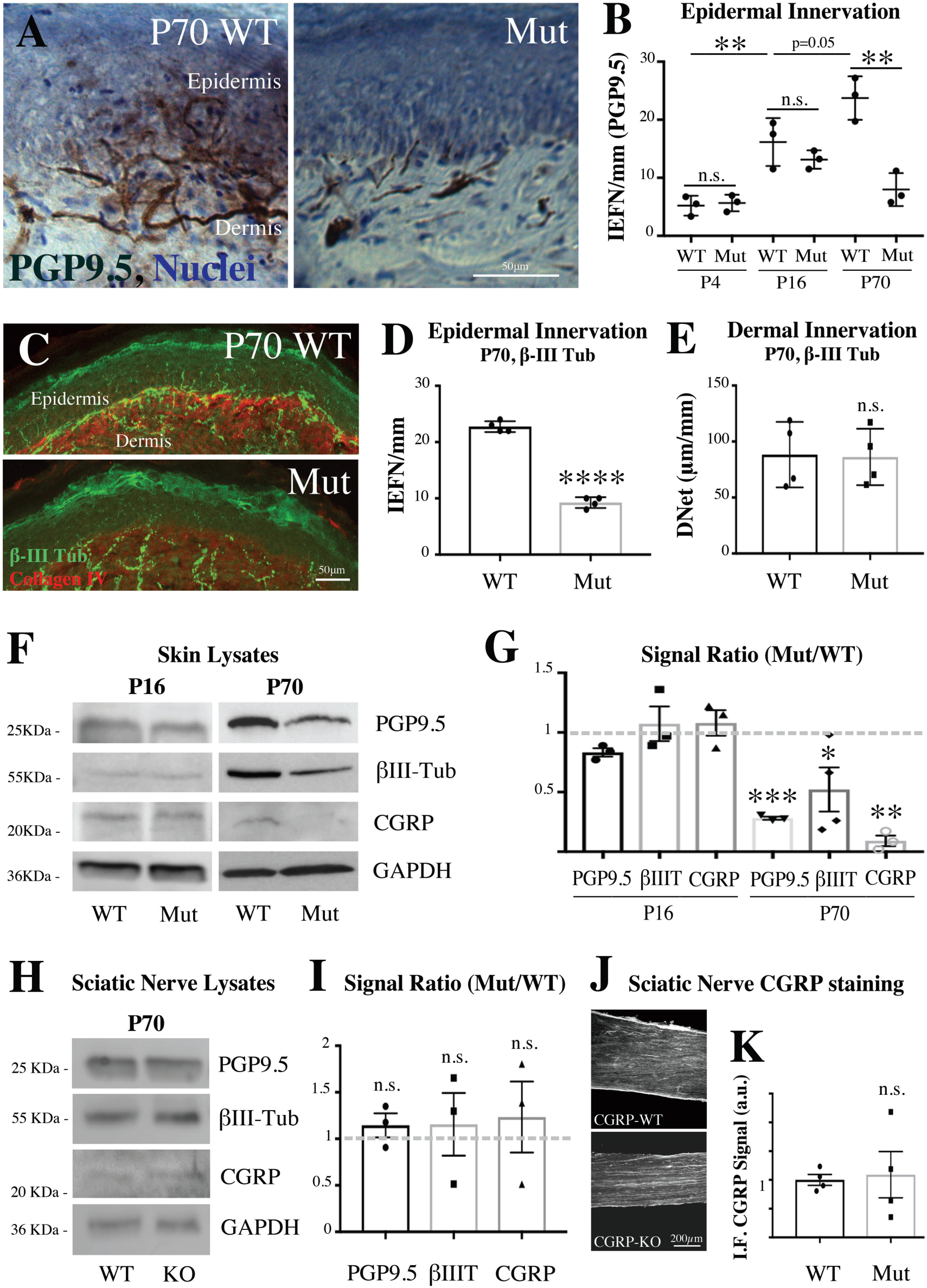
Cdkl5 is required for epidermal innervation. **A.** Immunohistochemical localization of Nerve Fibers in the Derm and in the Epiderm of 50 µm murine skin sections (P70), using PGP9.5 as specific axonal marker (20x magnification, Zeiss LSM-780 inverted confocal microscope). Sections were counterstained for nuclei in Mayer’s Heamatoxylin. **B**. The density of the intra-epidermal nerve fibers (IEFN/mm), crossing the derm-epidermal junction, was analyzed in WT and *Cdkl5^-/y^* mice at different developmental stages (P4, P16, P70). N:3, Mean with SD, Two way ANOVA, Sidak’s post-hoc, \*\**P<0.005*. **C**. Immunofluorescence of skin biopsies from P70 WT and *Cdkl5^-/y^* mice. Representative micrographs where collagen IV identifies the epidermal basal lamina and βIII-tubulin identifies intra-epidermal nerve fibers (20x magnification, Zeiss LSM-780 inverted confocal microscope). **D**. Individual IENF crossing the dermal–epidermal junction were analysed. N:4, Mean with SD, Student’s *t*-test, *\*\*\*\*P<0.0001.* **E**. The intra-dermal neuronal network (DNet, µm/mm) was measured by using Neuron-J software (magnification 20x). N:4, Mean with SD, Student’s *t*-test, *P>0.05*. **F**. Western blotting showing the expression of neuronal markers (PGP9.5, βIII-tubulin and CGRP) from skin lysates of P16 and P70 WT and *Cdkl5^-/y^* mice. **G**. The immunoblotting bands have been quantified by densitometry after normalization with GAPDH used as internal loading control. N: 3-4 independent experiments, Mean with SD, Two way ANOVA, Sidak’s post-hoc, \*\*\*\**P<0.0001 **P<0.005*. **H**. Western blotting showing the expression of PGP9.5, βIII-tubulin and CGRP from sciatic nerve lysates in WT and *Cdkl5^-/y^* mice. GAPDH was used as loading control. **I**. Densitometry of immunobloting bands showing no difference between P70-WT and *Cdkl5^-/y^* mice was performed after normalization to GAPDH. N: 3 independent experiments, Mean with SD, Student’s *t-*test, *P>0.05*. **J**. Representative CGRP immunofluorescence of sciatic nerve sections from P70-WT and *Cdkl5^-/y^* mice (10x, Nikon Eclipse-TE-2000U microscope). **K**. Graphs showing quantification of the immunofluorescence signal display no difference between WT and *Cdkl5^-/y^* mice. N: 3 mice per group, Mean with SD, Student’s *t-*test, *P>0.05*.

### Cdkl5 is required for TRPV1/CaMKII-dependent signalling and nociception in sensory neurons and in vivo

Since TRPV1 activity relies on CaMKII-dependent interaction and phosphorylation (*33, 34*), we hypothesized that TRPV1 signalling might require Cdkl5. Therefore we aimed to address whether Cdkl5 and CaMKII regulate TRPV1-dependent calcium signalling after stimulation with the well-established ligand capsaicin, which activates nociception by engaging with its receptor TRPV1 (*35–37*).

We used capsaicin as specific activator of TRPV1 and we measured calcium influx in iPS derived nociceptors to find that capsaicin-induced increase in calcium is impaired in neurons carrying CDKL5-mutations involving the kinase active domain (E55fsX74) and the TEY motif (G155fsX197) (Fig. 4C). Similarly, capsaicin-induced calcium responses were impaired in Cdkl5 mutant murine DRG explants and cultured neurons, where we established that overexpression of AAV9-CDKL5 rescued the defect in calcium response (Fig. 4D,G). When we blocked CaMKII signalling in cultured WT DRG neurons by using KN93, in contrast to its inert structural analog KN92, we observed, as expected, that KN93 inhibited the activation of TRPV1 after administration of capsaicin. Inhibition was similar to what observed in *Cdkl5* mutant cells, where the CaMKII inhibitor did not further decrease calcium levels (Fig. 4E). Delivery of KN93 after overexpression of CDKL5 in *Cdkl5* mutant DRG neurons blocked the rescue in calcium levels after capsaicin delivery (Fig. 4F). Together, these data confirm our initial hypothesis that CaMKII is required for Cdkl5-dependent calcium responses after capsaicin.

Finally, we hypothesized that capsaicin signalling would also be compromised in vivo leading to biochemical and behavioural impairments in *Cdkl5* deficient mice. In line with this hypothesis, we found that both the licking behaviour and cell signalling response to intradermal injection of capsaicin were impaired *in vivo* in *Cdkl5^-/y^* mutant mice and following conditional deletion of Cdkl5 specifically in sciatic DRG sensory neurons. In fact, the time spent licking the paw after injection of capsaicin (1µg) was significantly reduced both in *Cdkl5^-/y^* mice and in AAV5-CRE vs AAV5-GFP treated *Cdkl5*-floxed mice (Fig. 6A-C). Consistently, five minutes after capsaicin injection, pCaMKIIα was upregulated in DRG in WT but not in *Cdkl5^-/y^* mice, supporting a defective signalling response (Fig. 6D-F). We confirmed defective transmission of the stimulus in *Cdkl5^-/y^* mice also to the dorsal horn of the spinal cord by measuring pERK1/2 expression, typically activated by capsaicin-TRPV1 signalling in lamina I-II of the dorsal horn (*38*). Indeed, pERK1/2 was induced only in WT but not in *Cdkl5^-/y^* mice following intradermal injection of capsaicin (Fig. 6G-I) (*39, 40*).

**Figure 6.**
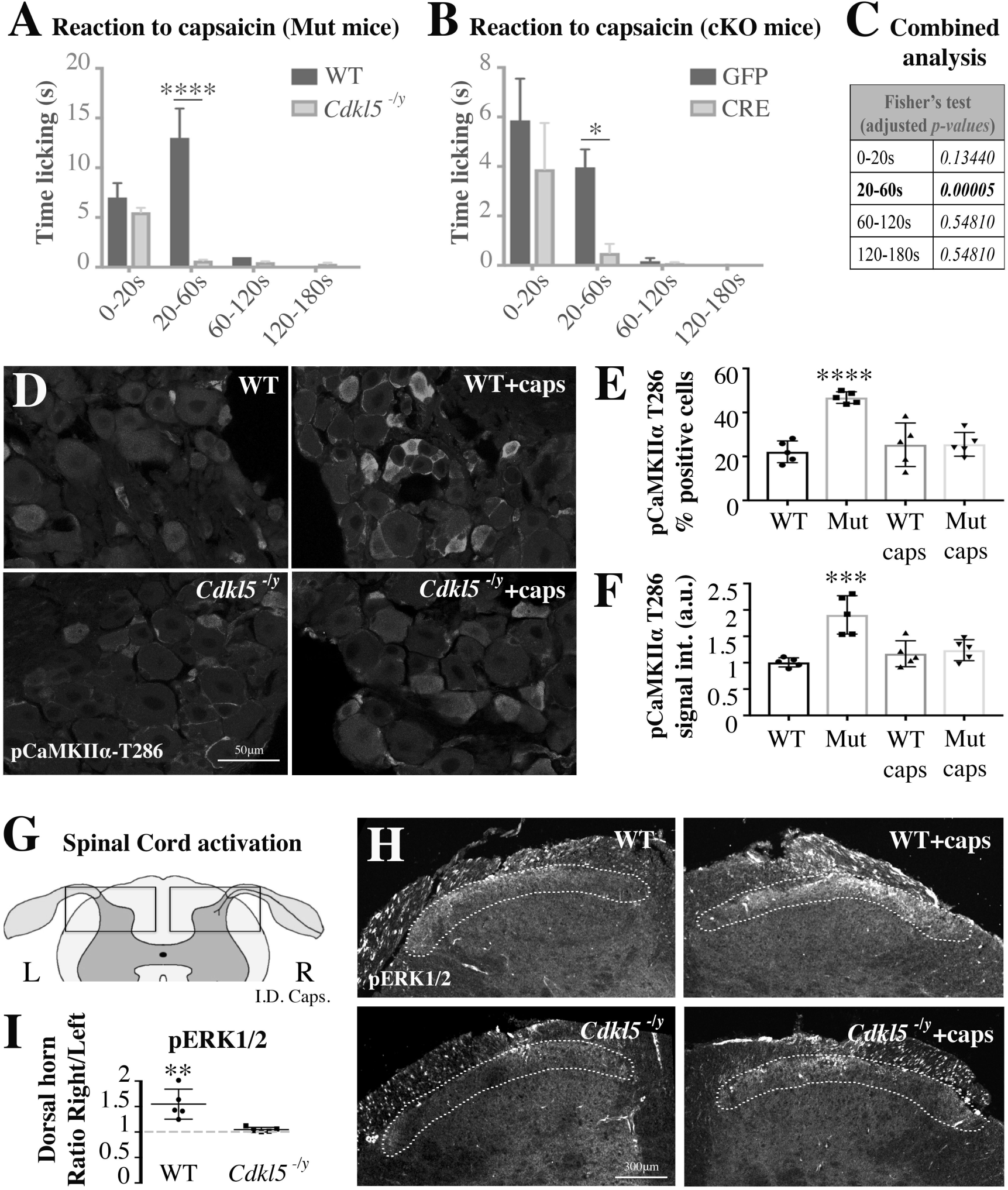
Cdkl5 deletion impairs capsaicin mediated nociceptive signalling and licking behaviour. **A,B**. Bar graphs showing the time spent licking the paw after the injection of 1µg of capsaicin in the posterior right paw vs vehicle injected in the left paw (A: WT vs *Cdkl5^-/y^*; B: GFP vs CRE AAV5 intra-sciatic injected *Cdkl5*-floxed mice). Time licking the paw is significantly reduced in *Cdkl5^-/y^* mice and in mice where *Cdkl5* was conditionally deleted (cKO) in DRGs. **A**. N: 5 mice each group, Mean with SEM, Two way ANOVA, Sidak’s post-hoc, \*\*\*\**P<0.0001.* **B**. N: 8 mice each group, Mean with SEM, Two way ANOVA, Sidak’s post-hoc, \**P<0.05.* **C**. Fisher’s combined probability test (false discovery rate, meta-analysis adjusted for multiple testing) of the *p-values* from the time courses in A and B (one statistical test for each time point). **D**. Representative confocal immunofluorescence images for phospho-T286-CaMKIIα in WT and *Cdkl5^-/y^* sciatic DRGs (63x, Leica TCS SP5 II). Tissue was fixed immediately after the capsaicin injection test. **E**, **F**. Bar graphs showing the percentage number of positive pCaMKIIα cells (**E**) and the pCaMKIIα average signal intensity in WT vs *Cdkl5^-/y^* DRGs-small cells after normalisation to WT (**F**). N: 5 mice per group, Mean with SD, One way ANOVA, Tukey’s post-hoc, \*\*\*\**P<0.0001* \*\*\**P<0.001*. **G**. Schematic illustrating a cross section of the lumbar spinal cord emphasizing the dorsal horns where nociceptive neurons make a synapse in the lamina I-II (L: left, R: right). **H**. Representative confocal immunofluorescence images for pERK1/2 in WT and *Cdkl5^-/y^* spinal dorsal horns (10x, Leica TCS SP5 II). Tissue was fixed five minutes after capsaicin injection in the posterior right paw vs vehicle injected in the left paw. Dotted lines indicate lamina I/II showing an increase in pERK1/2 signal of WT but not *Cdkl5^-/y^* mice. **I**. Graph showing lack of pERK1/2 activation (signal intensity) in spinal lamina I-II (right side vs left side) in *Cdkl5^-/y^* mice vs WT. N: 5 mice per group, Mean with SD, Student’s *t-*test, \*\**P<0.01*.

## Discussion

Our data reveal a previously unknown function of Cdkl5 in the regulation of primary nociception in the peripheral nervous system as shown by impaired nociception following conditional deletion of *Cdkl5* in DRG sensory neurons. Specifically, we found that Cdkl5 is required for CaMKII-dependent TRPV1 signalling and outgrowth of sensory neurons in both CDD murine and human neurons. *In vivo,* in a CDD animal model, this translates into impaired capsaicin-dependent nociceptive signalling and behavioural responses as well as in reduced epidermal innervation.

The anatomical, mechanistic and functional data all together support the anamnestic evidence that over 34% of CDKL5 patients have defective nociception. In a much smaller percentage of patients with enhanced pain perception, cortical hyperexcitability, which is a typical feature of these patients, might lead to dysfunctional central processing of nociception. This might be in line with disorders affecting cognition and neuronal activity in the central nervous system such as ASD and Rett syndrome (RTT), where it has been proposed that altered pain sensitivity is related to a dysfunctional mode of cortical pain processing, rather than to defective primary peripheral nociception (*41*) (*42–45*). CDD and RTT are distinct diseases caused by mutations in separate independent genes, *Cdkl5* and methyl CpG binding protein 2 (*MeCP2*) respectively. However, in vitro data suggested that the two might interact (*1, 46, 47*), although functional interaction in vivo remains unclear.

Recently, genes such as *MeCP2* and Ca2+/calmodulin-dependent protein kinase IIα (*CaMKIIα*), which have been implicated in the pathogenesis of ASD (*48–50*), and that are highly expressed in the CNS, have also been localized to the peripheral nervous system where they regulate sensory modalities (*51, 52*). However, a role in nociception had not been described so far.

Limitations of our work include the need for future investigation into potential additional Cdkl5-dependent mechanisms that might control nociception. Additionally, it is presently not known whether Cdkl5 plays a role in inflammatory or in traumatic painful conditions.

In summary, the importance of these findings lies on the (i) identification of a previously unrecognized regulatory mechanism for nociception that relies on Cdkl5, the (ii) discovery of impaired nociception as a previously uncharacterized symptom in CDD and (iii) the usefulness of nociception as therapeutic outcome measure in animal models of CDD. Since animal models of CDKL5 deficiency do not develop seizures, monitoring pain responses during therapeutic interventions might be a unique opportunity to test novel disease-modifying treatment in pre-clinical before clinical settings. Lastly, our data suggest that gene therapy or other interventions in CDKL5 patients should not only be directed to the brain but also to dorsal root ganglia to restore nociceptive input in patients with impaired nociception.

## Research Design and Methods

### Questionnaire

Ethics approval was obtained from the Human Research Ethics Committee, University of Western Australia.

### Mice

All animal procedures were approved by Imperial College London ethic committee, and were performed in accordance with the UK Animals Scientific Procedures Act (1986). C57Bl/6 background mice lacking *Cdkl5* exon 6 (*4*) and Cdkl5 floxed C57Bl/6 background mice containing loxP sites flanking *Cdkl5* exon 4 (*12*) and wild-type littermates or C57Bl/6 (Harlan, UK) mice ranging from day-4 to day-70 of age were used for all experiments. Mice were anaesthetized with isoflurane (3% induction, 2% maintenance) and buprenorphine (0.1  mg  kg^−1^) and carprophen (5  mg  kg^−1^) were administered peri-operatively as analgesic. The experimenter was blind to the genotype of animals and to treatment given to each animal.

### Epidermal innervation

Following euthanasia, the external skin of the mouse foot was sterilized with ethanol and a skin punch, 1 mm in diameter, was introduced perpendicularly to the surface of pad skin, rotated and advanced until reaching 1-2 mm depth (*53*). The specimens were immediately fixed in 3% paraformaldehyde, 75mM Lysine hydrochloride (Sigma L5501), 2.2mg/ml Sodium Metaperiodate (Sigma S1878) for up to 24 hours at 4 °C, and then kept in a cryoprotective solution overnight. After fixation, the tissue was serially cut (sections of 50 µm in thickness, perpendicular to the dermis) using a freezing microtome. The slices were then treated with blocking solution for two hours at RT (PBS – Triton X-100 0.3% - Donkey Serum 10%) and exposed to different primary Abs. βIII-Tubulin (Abcam ab-18207) and Collagen IV (Sigma C1926) were the markers used to assess, respectively, the density of IENF and the basal lamina. Using confocal microscopy (Leica TCS SP5 II or Zeiss LSM-780 inverted confocal microscope), individual IENF crossing the dermal–epidermal junction were counted and the intraepidermal network length was analyzed by using Neuron-J software (magnification X20) in at least three non-consecutive sections (*54*).

For anti-PGP9.5 immunohistochemistry, sections were immunostained by using Vectastain ABC kit (PK-4001) with minor adjustments. Endogenous peroxidase was blocked by incubation in PBS/0.1% Tx100 containing 0.3% hydrogen peroxide for 10 min. After 2x PBS washes, sections were blocked in normal blocking serum for 20min and incubated with primary Ab (anti-PGP 9.5, Ultraclone, 1:20000, gift from Prof. Praveen Anand) overnight at R.T. After washing, sites of primary Ab binding were revealed by biotinylated secondary anti-rabbit incubation followed by avidin–biotin peroxidase reaction, using Impact DAB as substrate (SK-4105). Sections were counterstained for nuclei in Mayer’s Heamatoxylin, dehydrated with serial incubations of ethanol: 70% (1x, 2min); 90% (1x, 2min); 100% (2x, 5min), and mounted in xylene-based mounting medium.

### Murine DRG, Spinal Cord and Sciatic Nerve Immunohistochemistry

After euthanasia, the tissues were removed and fixed in 4%-PFA-PBS for up to 24 h at 4 °C, and then kept in a cryoprotective solution overnight. After fixation, the samples were cut (thickness 15 µm) using a freezing microtome. Immunohistochemistry on tissue sections was performed according to standard procedures. For all antibodies used, the samples were blocked for 1  h with 10% Donkey Serum - 0.3% PBS–Triton-X100, and then incubated with Cdkl5 (rabbit, Sigma HPA002847, 1:100), CGRP (mouse, abcam ab81887, 1:100), IB4 (Thermo-Fisher I21411Alexa Fluor™ 488 Conjugate, 1:250), Parvalbumin (mouse, abcam ab64555, 1:100), TRPV1 (guinea pig, Thermo-Fisher PA1-29770, 1:100), CaMKIIα (mouse, Thermo-Fisher MA1-048, 1:100), phospho-Thr286-CaMKIIα (mouse, Thermo-Fisher MA1-047, 1:100), phosphor-Thr202/Tyr204-Erk1/2 (rabbit, Cell Signaling #9101, 1:100); βIII-Tubulin (rabbit, abcam ab18207, 1:500) antibodies at 4  °C overnight. Subsequently, samples were incubated with AlexaFluor-conjugated goat secondary antibodies according to standard protocols (Invitrogen). All tissue sections were counterstained with Hoechst (Molecular Probes).

### Differentiation of iPSC Derived Nociceptors

iPSC lines from four CDKL5 patients, carrying different mutations (E55fsX74, G155fsX197, Q347X, S855X; special gift from Prof. N. Mazarakis, Imperial College London, and Dr. D. Millar, Cardiff University), were cultured in Matrigel coated wells (Corning Membrane Matrix 354234) and mTeSR^TM^1 medium (STEMCELL Technologies) supplemented with the ROCK inhibitor Y27632 10µM (STEMCELL Technologies). When a confluency of 60-70% was reached, we started the differentiation by using five inhibitors for 10 days (LDN193189 100nM, SB431542 10µM, CHIR99021 3µM, SU5402 10µM, DAPT 10µM; STEMCELL Technologies) in hESC medium (DMEM-F12, knockout serum replacement 20% (Gibco), L-Glut 1mM, MEM non essential aminoacid solution (M7145 Sigma)) (*55*). The cells were then grown in Neuronal Growth Medium (DMEM-F12, HI-FBS 10%) supplemented with neurotrophins (BDNF 10ng/ml, NGF 10ng/ml, NT3 10ng/ml, GDNF 10ng/ml; STEMCELL Technologies) and ascorbic acid (35ng/ml; 72132 STEMCELL Technologies) for 4 days and treated with mitomycin C (1µg/ml; 73272 STEMCELL Technologies) for 3 hours. Lastly, the cells were grown for 30 days and replated 1 day before the calcium imaging experiment. The expression of neuronal markers (βIII-Tubulin and CGRP for nociceptors) was monitored at days 0, 7 and 12, as well as at day 42, when we confirmed their sensitivity to 50mM KCl and 500ng/ml capsaicin.

Cells were fixed with 4% PFA – 4% sucrose and the immunocytochemistry was performed by incubating fixed cells with βIII-tubulin (Abcam ab18207) and CGRP (Abcam ab81887) antibodies at 4  °C overnight. This was followed by incubation with AlexaFluor-conjugated goat secondary antibodies according to standard protocols (Invitrogen). All cells were counterstained with Hoechst (Molecular Probes).

### Western Blot analysis of skin, brain, DRG and sciatic nerve lysates

Approximately 10-30 µg of protein extracts (lysis buffer: 50 mM Tris-HCl, pH 7.4, 150mM NaCl, Nonidet P-40 1%, 1 mM EDTA, 0.5 mM DTT, protein inhibitor cocktail Sigma, 1 mM PMSF; lysed for 30  min on ice followed by 15  min centrifugation at 4  °C) from brain, DRGs, sciatic nerve and skin were heated at 95°C and separated by 8% SDS–polyacrylamide gel electrophoresis (PAGE) gels (37.5:1 acrylamide:bis-acrylamide) and transferred to nitrocellulose membranes for 2  h. Before gel loading, protein concentration was quantified using Pierce BCA Protein Assay Kit (ThermoScientific). Membranes were blocked with 5% BSA or milk for 1  h at room temperature and incubated with Cdkl5 (Sigma HPA002847, 1:1000), CaMKIIα (avivasysbio Ab286 OAECOO614, 1:1000), CGRP (abcam ab-189786, 1:1000), PGP9.5 (abcam ab-8189, 1:1000), βIII-tubulin (Abcam ab-18207, 1:1000), Gapdh (Cell Signaling #2118) antibodies at 4  °C overnight. Following HRP-linked secondary antibody (GE Healthcare) incubation for 1  h at room temperature, membranes were developed with ECL substrate (GE Healthcare) (ThermoScientific) and the bands detected with a CCD camera (Syngene, GeneGnome XRQ) and analyzed, after subtraction of the background, by using the ImageJ software.

### RNA isolation and RT-PCR Analysis

Total RNA was isolated using the RNeasy Mini Kit (Qiagen) according to the manufacturer’s instructions. Contaminating DNA was removed with DNaseI (Invitrogen 1734510), and the obtained RNA was quantified with a NanoDrop UV Visible Spectrophotometer; the quality of the RNA was assessed through agarose gel electrophoresis. cDNA was synthetized from 200 ng of RNA using the SuperScript First Strand reverse transcriptase kit (Invitrogen 1933910) as indicated by the manufacturer. The RT-PCR was performed in 20 µl using 50 ng of cDNA and the following cycling parameters: *Cdkl5-Ex6* and *Gapdh*, 35 cycles, 95 °C 1m, 54 °C 30s, 72 °C 1m. The forward (F) and reverse (R) primers were:

m*Cdkl5-*Ex6-Fw, 5’-GGAGACGACCTTACGAGAGC

m*Cdkl5-*Ex6-Rv, 5’-GGACGATGTCGTTCTTGTGG (PCR product: 236bp)

m*Gapdh* F, 5’-AAGGTCGGTGTGAACGGATTTG

m*Gapdh* R, 5’-GCAGTGATGGCATGGACTGTG (PCR prduct: 536bp).

### Injection of viral vectors (AAV5-GFP; AAV5-Cre-GFP) into the sciatic nerve

2.5  µl of each viral vector (AAV5-GFP SignaGen SL100819 or AAV5Cre-GFP SignaGen SL100821, titer 3.06X10^13^ GC/ml) was injected into the sciatic nerve of adult *Cdkl5*-floxed mice (*12*) with a Hamilton syringe and Hamilton needle (NDL small RN ga34/15mm/pst45°). After 2 months, the animals were exposed to specific behavioural tests and then sacrificed, to evaluate the silencing of *Cdkl5*gene in sciatic DRGs by WB.

### Mechanical Allodynia Test

Mechanical allodynia was quantified by measuring the hind paw withdrawal response to von Frey filament stimulation. Animals were placed in methacrylate cylinders (20 cm high, 9 cm diameter) with a wire grid bottom through which the von Frey filaments (North Coast Medical,

Inc., San Jose, CA) with a bending force in the range of 0.16-8g were applied by using a modified version of the up–down paradigm, as previously reported by Chaplan *et al*. The filament of 0.16g was used first and the 8-g filament was used as a cut-off. Then, the strength of the next filament was decreased or increased according to the response. The threshold of response was calculated using a regression curve generated based on the sequence of filament number (numbered from 1 to 9) versus log(10) of their strength in gr, an interpolation of the number of filament plus or minus a correction factor of 0.5 according to the response was then used to calculate the threshold of response in gr. Clear paw withdrawal, shaking, or licking of the paw was considered as a nociceptive-like response. Both hind paws were tested and averaged. Animals were allowed to habituate for 1 h before testing in order to allow an appropriate behavioural immobility.

### Hargreaves’ Test (Plantar Test)

*Thermal hyperalgesia* was assessed as previously reported by Hargreaves *et al*. Paw withdrawal latency in response to radiant heat was measured using the plantar test apparatus (Ugo Basile, Varese, Italy). Briefly, the mice were placed in methacrylate cylinders (20 cm high × 9 cm diameter) positioned on a glass surface. The heat source was positioned under the plantar surface of the hind paw and activated with a radiant light beam, intensity was chosen in preliminary studies to give baseline latencies from 8 to 9 s in control mice. A cut-off time of 15s was used to prevent tissue damage in the absence of response. The mean paw withdrawal latencies from both hind paws were determined from the average of five separate trials, taken at 2-min intervals to prevent thermal sensitization and behavioural disturbances. Animals were habituated to the environment for 1 h before the experiment to become quiet and to allow testing (*56*).

### Grid walk

Mice were allowed to run the grid walk (50  ×  5  cm plastic grid (1  ×  1  cm) placed between two vertical 40-cm-high wood blocks) three times per session. The total number of steps and missteps per run for each hindpaw was analyzed by a blinded investigator.

### Adhesive removal

Small adhesive stimuli (6-mm round adhesive labels) were placed on both hindpaws of the mouse, the time to make first contact with both forepaws, as well as the time to completely remove the adhesive were recorded for each paw. Each mouse underwent three trials. All testing was performed by a blinded investigator, after a period of acclimatization.

### Cdkl5 Immunoprecipitation

The cortex of three mice (P65) was removed and immediately lysed in lysis buffer (Tris-HCl 7.5 20mM, NaCl 150mM, EDTA 1mM, EGTA 1mM, Triton X-100 0.1%, PIC Sigma P8340-1x, 1mM DTT, PhosSTOP Sigma P0044-1x) on ice (glass pestle Thermo-Fisher FB56679) and centrifuged (14000g 15m). The protein concentration was determined by the Bradford method. For each sample, 1mg of protein was treated for 30m with 500U Benzonase (E1014 Sigma) at 37°C, then exposed (overnight wheel, 4°C) to 1µg of Cdkl5-Ab (Sigma HPA002847) or 1µg Rabbit IgG (Jackson 011000003 lot95014). Magnetic beads (40µl 50% Dynabeads Protein G 10003D, Life Technologies) were subsequently added to the samples, and the mixture was further incubated on the wheel for 2hrs at 4°C. The beads were washed three times (3×20m washes with lysis buffer, wheel 4°C) and proteins were eluted by boiling for 10 min in NuPage Sample Reducing Agent - DTT (30µl, Thermo-Fisher NP0009; 80°C) and loaded on a 8% SDS-PAGE or analyzed by Mass Spectrometry.

### Mass Spectrometry Sample Preparation

Samples were boiled at 70°C for 10 minutes in 1x NuPAGE LDS Sample Buffer (Life technologies) containing 100mM DTT and separated on a 10% NuPAGE Bis-Tris gel (Life technologies) for 10 at 180V in MES running buffer (Life technologies). After fixation in 7% acetic acid containing 40% methanol and subsequently staining for 30 minutes using Colloidal Blue staining kit (Life technologies) protein lane was excised from the gel, chopped and destained (50% ethanol in 25 mM NH4HCO3) for 15 minutes rotating at room temperature and dehydrated for 10 minutes rotating in 100% acetonitrile. Vacuum dried samples were rehydrated and reduced for 60 minutes in reduction buffer (10mM DTT in 50mM NH4HCO3 pH 8.0) at 56°C and subsequently alkylated in 50 mM iodoacetamide in 50mM NH4HCO3 pH 8.0 for 45 minutes at room temperature in the dark. Dehydrated and vacuum dried samples were trypsin digested (1µg trypsin/sample in 50mM Triethylammonium bicarbonate buffer pH 8.0) at 37°C overnight. Stepwise peptide extraction was done as follows: twice extraction solution (30% acetonitrile) and 100% acetonitrile for 15 minutes at 25°C shaking at 1400 rpm. Reductive methylation for quantification was performed as described in Hsu et al. (2003) (*57*). For Cdkl5 immunoprecipitation, peptides were labelled LIGHT (DimethLys-0 and DimethNter0) and control IgG immunoprecipitation peptides labelled HEAVY (DimethLys-4 and DimethNter4). Afterwards they were mixed 1:1 and further processed. In the so called “reverse” experiment Cdkl5 IP peptides were labelled HEAVY and control IgG IP peptides were labelled LIGHT. We performed Cdkl5 IP experiments in duplicates for each of the three different animals resulting in n=6. After purification and desalted using C18 stage tips (Empore) (*58*) 3.5 µL peptides were loaded and separated on C18 column (New Objective) with 75 µm inner diameter self-packed with 1.9µm Reprosil beads (Dr. Maisch) which was mounted to an EasyLC1000 HPLC (Thermo-Fisher Scientific).

### Mass Spectrometry Measurement and Data Analysis

Reversed-phase chromatography gradient (Buffer A: 0.1% formic acid, Buffer B: 80% acetonitrile and 0.1% formic acid, Gradient: 0-67 min 0-22% Buffer B, 67-88 min 22-40% Buffer B, 89-92 min 40-95% Buffer B) was applied and peptides eluted and directly sprayed into a Q Exactive Plus mass spectrometer from Thermo-Fisher Scientific operating in positive scan mode with a full scan resolution of 70,000; AGC target 3×10∧6; max IT = 20ms; Scan range 300 - 1650 m/z and a Top10 MSMS method. Normalized collision energy was set to 25 and MSMS scan mode operated with resolution of 17,000; AGC target 1×10∧5; max IT = 120 ms. Triggered MSMS masses were excluded dynamically for 20 seconds. Database search was performed using MaxQuant Version 1.5.2.8 (*59*) against Mus Musculus Uniprot database (downloaded 8th January 2018; 53819 entries) with Trypsin/P as digestion enzyme allowing 2 missed cleavages. As settings the following was applied: variable modification: Acetyl (Protein N-term); Oxidation (M), fixed modifications: Carbamidomethyl (C), FDR of 1% on peptide and protein level was applied. As light label: DimethylLys0 and DimethylNter0 and heavy label: DimethylLys4 and DimethylNter4 were set with max. 3 labeled amino acids. Proteins with at least two unique peptides were considered as identified. Proteins matching reverse database or common contamination list as well as proteins with peptides only identified by peptides with modification were filtered out. Statistical calculation and visual presentation was done in R version 3.5.1 (released 2^nd^ July 2018; R Core Team (2016) R: A language and environment for statistical computing R: Foundation for Statistical Computing, Vienna, Austria. URL https://www.R-project.org/). Protein ratios of each immunoprecipitation experiment (n=6) were divided by median to normalized the ratios. The new obtained median was 1. All “reverse” replicates were inverted to obtain, positive values for IP enrichment. Log2 transformed median normalized ratios were plotted as box plot (function “ggplot“, package “ggplot2“). For each individual protein depicted in the box plot a parametric Two-tailed tests was performed (function “compare_means”, package “ggpubr”) against all other measured proteins. Significance was calculated and indicated (\**P<0.05* \*\**P<0.01* \*\*\**P<0.001* \*\*\*\**P<0.0001*). Further packages used: “openxlsx”, “dplyr”, “reshape”. These studies were performed in the proteomics facility in Mainz, Germany.

### DuoLink Proximal ligation Assay

After blocking DRGs slices (50µm of thickness), from P65 C57Bl/6 *Cdkl5* null mice and wild-type littermates with Duolink Blocking Solution, primary antibodies raised in two different species, specific for the two proteins of interest (rabbit-Cdkl5 Sigma HPA002847 and mouse-CaMKIIα Thermo-Fisher MA1-048), were added to the tissue samples (1:100) and incubated in a humidity chamber overnight at 4°C. After washing, according to the manufacturer protocol (PLA; Sigma DUO92101), the samples were then incubated with secondary antibodies coupled to oligonucleotides (proximity probes) for 1 hour at 37°C and exposed to the ligase for 30 minutes at 37°C; the DNA polymerase was lastly added for 100 minutes at 37°C. The slides were washed thereafter, mounted and analyzed at the Leica TCS SP5 II confocal microscope (63x magnification; 6 non-consecutive 38µm-sections from the entire DRG).

### DRG culture

Adult DRGs were dissected and collected in Hank’s Balanced Salt Solution (HBSS) on ice and digested according to standard procedures. Cultures were maintained in medium containing B27 and penicillin–streptomycin in DMEM:F12. Cells were plated on coated glass coverslips (0.01  mg  ml^−1^PDL) for 48-72  h, and fixed with 4% PFA–4% sucrose. Immunocytochemistry was performed by incubating fixed cells with Cdkl5 (Sigma HPA002847), βIII-tubulin (abcam ab18207), CaMKIIα (Thermo-Fisher MA1-048), (abcam ab81887), Parvalbumin (abcam ab555), GFP (Thermo-Fisher A-6455), HA (abcam ab18181) antibodies at 4  °C overnight. This was followed by incubation with AlexaFluor-conjugated goat secondary antibodies according to standard protocols (Invitrogen). All cells were counterstained with Hoechst (Molecular Probes).

### DRG explants

DRGs from adult WT and *Cdkl5* null mice were isolated, plated on a culture dish and incubated in Ca(2+) dye Fluo-4AM (Thermo-Fisher F14201 494/506nm; 1µM) in HBSS (with 20mM Hepes pH7.4 and 1µl/ml pluronic F127 P3000MP, Life Technologies) for 30m at 37°C. After washing in HBSS-Hepes, the DRGs were placed on the surface of polystyrene-plates, perfused with a small amount of “basal medium” (2ml/m: NaCl 160mM, KCl 2.5mM, CaCl_2_ 1mM, MgCl_2_ 2mM, Hepes pH7.4 10mM, glucose 10mM; 37°C) and stimulated for 20s with capsaicin (Tocris0462 500ng/ml). Live-imaging was acquired by Nikon Eclipse-TE-2000U microscope (20x magnification) and analyzed by ImageJ (Fluo-4 signalling: increased fluorescence emission at 506nm).

### DRG culture infection and treatments

WT and Cdkl5 null DRG-neurons seeded at a density of 5,000 cells/cm^2^ were infected with AAV9 viral particles (AAV9-Ctrl-ss-pTR-CBh-GFP-stuffer; AAV9-ss-pTR-CBh-HA-CDKL5_107_) 24h after plating (100,000 virus genomes (VG)/cell) (*60*). Microscopic analysis was carried out 24 hrs post transduction with a Nikon Eclipse-TE-2000U microscope (20x magnification, bright-field phase contrast or GFP-488nm). 24, 48 and 72 hrs after plating, the neurons were treated with KN92, KN93 (0.5µM, Tocris 4130 and 1278) or DMSO alone (*61*). 72 hours after the treatment or the infection, the respective efficacy of KN93 and AAV9 particles was investigated (levels of phospho-CaMKIIα and Cdkl5-protein expression).

Cdkl5-protein Expression Analysis: 72 hrs after the treatment, the medium was removed and the cultures were scraped into RIPA buffer (50 mM Tris-HCl pH 7.4, 150 mM NaCl, 1% NP-40, 0.5% deoxycholate, protease and phosphatase inhibitors cocktails Sigma) and centrifuged at 14000g for 15 min at 4°C to remove cell debris. The supernatant was transferred to a new tube, mixed with Laemmli sample buffer (2x), boiled at 95°C for 5 min and loaded on SDS-PAGE (incubation with Cdkl5 (Sigma HPA002847, 1:1000) and βIII-tubulin (Abcam ab-18207, 1:1000) antibodies).

Analysis of the levels of Phospho-CaMKIIα: 72 hrs after the treatment, the wells were fixed and processed with anti-phospho-Thr286-CaMKIIα (mouse, Thermo-Fisher MA1-047, 1:100) and beta-III-tubulin (rabbit, abcam ab18207, 1:500) antibodies, as previously described.

### Neurite outgrowth analysis

Immunofluorescence was detected using a Nikon Eclipse-TE-2000U microscope at 20× magnification using a CDD camera (scMOS camera QImaging). Five fields per coverslip were included in the analysis. At least 30 cells per coverslip were measured. Total neurite length was measured and normalized to the number of cells. All analyses were performed blinded. Neurite analysis and measurements were performed using the Neuron-J plugin for ImageJ.

### Image analysis for immunohistochemistry and immunocytochemistry

Photomicrographs were taken with Nikon Eclipse-TE-2000U microscope at 20× magnification. Alternatively, for confocal imaging, a Leica TCS SP5 II or a Zeiss LSM-780 inverted microscope (*z*-stacks, slice spacing 1.65µm) were used at 20× or 63x magnification. DRG, spinal cord and sciatic nerve micrographs were processed with the software ImageJ: a constant fluorescence intensity threshold was set across each cell (or area) in control-tissues. On the basis of this threshold, for each cell (or area) in the samples, the intensity of pixels was calculated in each channel. This was done in triplicate and the investigator was blinded to the experimental group.

### Single-cell calcium imaging

We assessed changes in the intracellular Ca2+ concentration ([Ca^2+^]_i_) using conventional ratiometric imaging in cultured DRG neurons isolated from WT and CDKL5^-/y^ mice. Briefly, after loading with 1µM Fura-2 (Invitrogen), the cells were exposed to capsaicin (500ng/ml) for 20 seconds and then to 50mM KCl (20 seconds) in (in mM) NaCl 160, KCl 2.5, CaCl_2_ 1, MgCl_2_ 2, Hepes 10, glucose 10 (pH7.4; 37°C) with a flow rate of ∼2ml/min. Recordings, with 355nM and 380nM excitation and at 0.5Hz, were done with the WinFluor (J Dempster, Strathclyde University), and analysed using the Clampfit (Invitrogen), software packages. Measurements were repeated at least in 3 independent cultures with analyzing ∼100 (50 WT vs 50 CDKL5^-/y^) cells in each. Responses of CDKL5^-/y^ neurons were normalized to those of WT cells (*62*).

### Intradermal injection of capsaicin

Capsaicin (10 µl) [0.1 g/ml^−1^ 100% ethanol; 1:1000 in PBS, 0.5% Tween 80, 10% ethanol] or vehicle (PBS, 0.5% Tween 80, 10% ethanol) was administered via intraplantar injection (*63*) to adult WT and *Cdkl5* mutant mice or to adult *Cdkl5*-Floxed mice treated, two months before, with AAV5-GFP or AAV5-GFP-CRE (intra-sciatic injection). After capsaicin injection, the licking time was measured during the first 3 minutes. Finally, WT vs *Cdkl5* null adult mice were sacrificed to study the activation of the peripheral pain pathways in fixed sciatic DRGs and spinal cord. Immunohistochemistry on DRG and spinal cord sections was performed by using anti-phospho-Thr286-CaMKIIα (mouse, Thermo-Fisher MA1-047, 1:100), or anti-phospho-Thr202/Tyr204-Erk1/2 (rabbit, Cell Signaling #9101, 1:100).

### HEK-293 Cell Transfection, CDKL5-Immunoprecipitation and Kinase Assay

HEK-293 cells were cultured on D-Poly-lysine in 60mm plates (DMEM Gibco, 0.7% glucose, 10% Fetal Bovine Serum, 200mM glutamine with 1% Penicillin/Streptomycin solution). After reaching 85% confluency, the cells were co-transfected with Lipofectamine vesicles (Thermo-Fisher L3000001 Lipofectamine-3000) containing two target DNA: pcDNA5-FRT/TO-C-3FLAG expressing rat-CAMKIIα and human-CDKL5 (or, alternatively, human-CDKL5-K42R); the control cells were treated with only Lipofectamine. The target plasmids (1ug per sample) and 15 µl Lipofectamine were added to the transfection medium Opti-MEM Gibco, left at room temperature for 15 minutes, to enable the transfer of plasmids into the vesicles, and then added to the HEK cells (incubation: 4 hours at 37°C). Finally, the transfection medium was replaced with definitive medium for three days. The cells were collected directly in 1ml of Lysis Buffer (20mM pH 7.4 Hepes, 150mM NaCl, 0.1% NP40, PIC Sigma P8340-1x, 1mM DTT, PhosSTOP Sigma P0044-1x), sonicated (1 min) and then centrifugated at 10000g (5 min, 4°C). The protein concentration of the supernatant was determined by the Bradford method and the CDKL5 Immunoprecipitation was performed as described (“Cdkl5 Immunoprecipitation”). To confirm the overexpression of the kinases, the 5% of the supernatant was directly loaded on SDS-PAGE. The kinase activity of CDKL5 was simultaneously verified using the supernatant of the lysate from plates overexpressing CDKL5 or its inactive form CDKL5-K42 (Kinase-Dead), exposed to Anti-Flag Beads (Sigma M2 Magnetic Beads; 2 hours, wheel 4°C). The beads were then washed three times (wheel 4°C, 20min) in Lysis Buffer, resuspended in Kinase Buffer-1x (Thermo-Fisher PV3189) and added to a mixture of 32P-ATP (1mM) and cold-ATP (1mM). After incubation (30 minutes at 30°C), the elution was performed by using Laemmli 6x (30 µl, 70°C, 3 min) and the samples were loaded on SDS-PAGE. The gel, fixed in methanol 2% (1 hour, RT), was dried (80°C, 2 hours) and exposed to an X-ray film for two days.

### Statistics and reproducibility

Data are plotted as mean  ±  S.D. All experiments were performed in triplicate unless specified. Asterisks indicate a significant difference analyzed by ANOVA with Tukey or Sidak post hoc test, Student’s *t*-test as indicated for normal distributions, parametric Two-tailed test and Fisher’s combined probability test. All data analysis was performed blinded to the experimental group.

## Acknowledgments

This work was supported by start-up funds from the Division of Brain Sciences, BRC funds, Imperial College London (SDG); Wings for Life (SDG); The Rosetrees Trust (SDG); the NIHR Imperial Biomedical Research Centre (SDG); International Foundation for CDKL5 Research (HL, JD, KW) and NHMRC (HL). We would like to thank Dr. Anja Freiwald for assistance with the proteomics data analysis (Proteomics facility in Mainz) and Dr. D. Millar, Cardiff University for providing iPS cells.

---

**Supplementary Figure 1.**
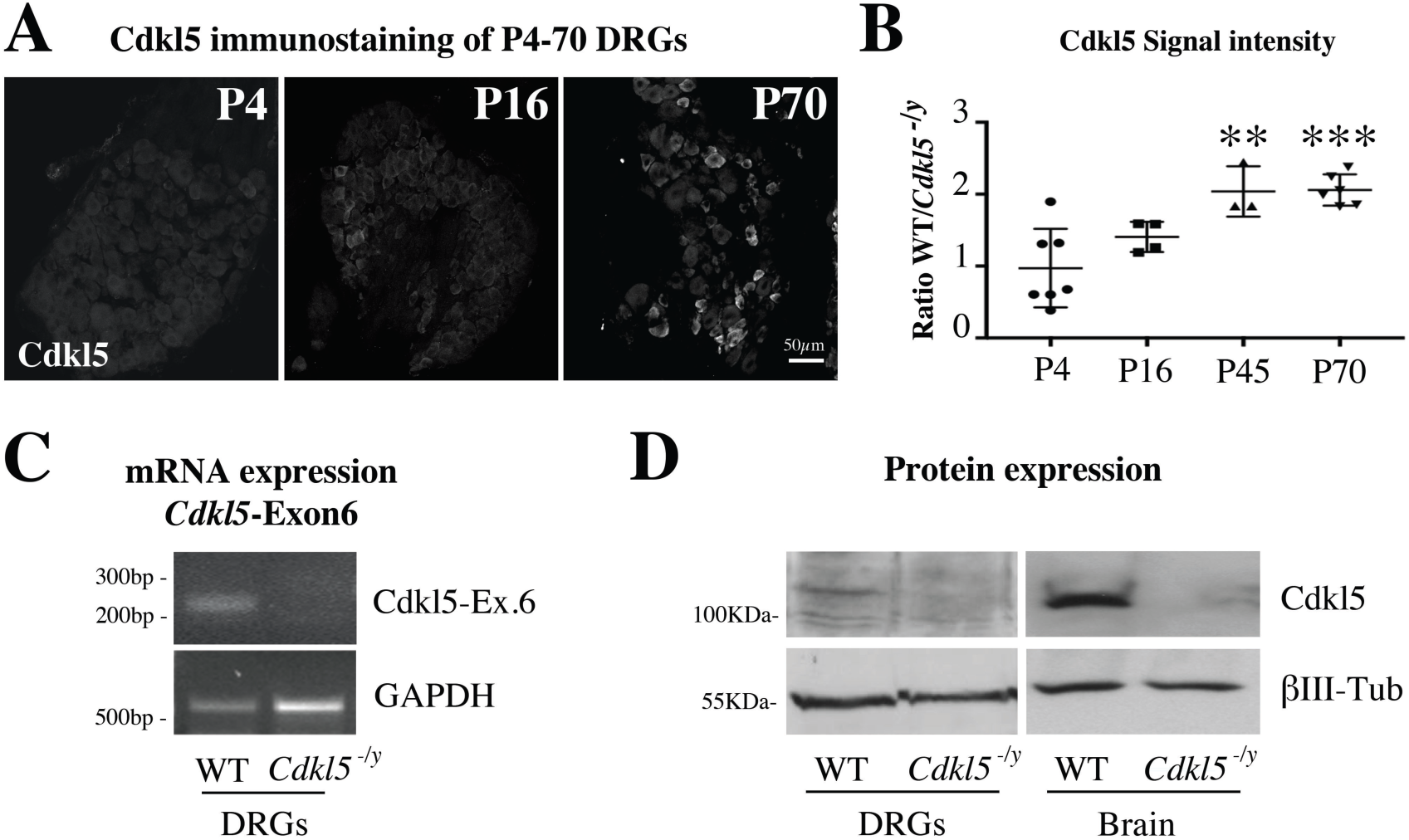
**A**. Representative images of immunostaining for Cdkl5 from WT P4, P16 and P70 DRGs (20x, Zeiss LSM-780 inverted confocal microscope) showing a clear positive signal in adult mice only. **B**. Graph showing quantification of the immunofluorescence signal displaying age-dependent expression levels (P4, P16, P45, P70). Ratio of the Cdkl5 signal intensity was calculated between WT and *Cdkl5^-/y^* DRGs to subtract the background signal. N: 3-7 mice per group, Mean with SD, One way ANOVA, Tukey’s post-hoc, \*\*\**P<0.001 **P<0.005***. C**. RT-PCR from WT and *Cdkl5^-/y^* adult DRG showing deletion of exon 6 in *Cdkl5^-/y^* mice. GAPDH has been used as control. **D**. Western blotting showing the expression of Cdkl5 in WT that is absent in *Cdkl5^-/y^* DRG and brain extracts. βIII-tubulin has been used as loading control.

**Supplementary Figure 2.**
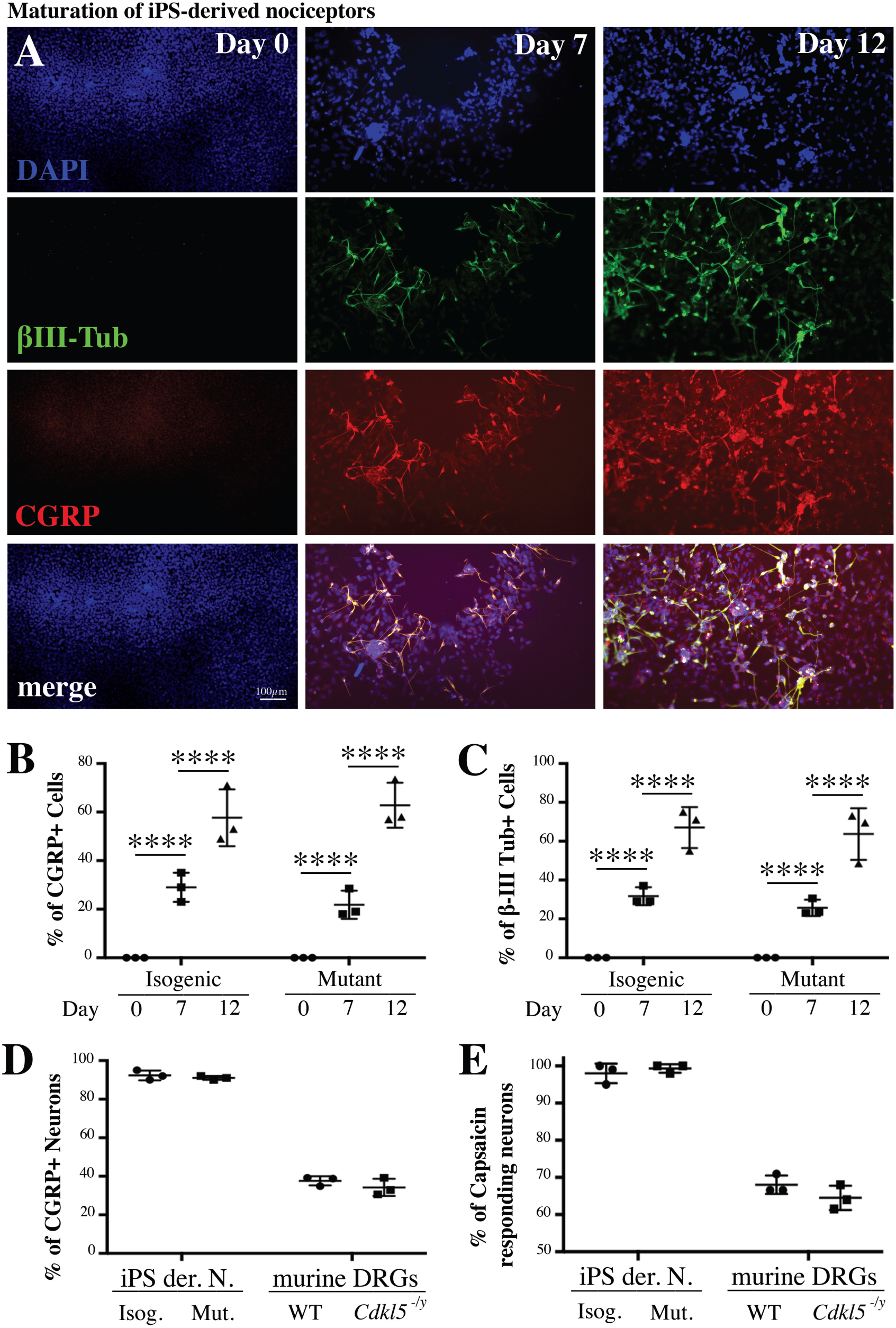
**A**. Representative immunofluorescence images for βIII-tubulin and CGRP (10x, Nikon Eclipse-TE-2000U microscope) from isogenic iPS at different stages of differentiation into nociceptors (day 0, day 7 and 12). **B, C**. Graphs showing the percentage of CGRP and βIII-tubulin positive cells at the three different maturation stages (DAPI staining was used to establish the total number of cells). N: 3 independent experiments (E55fsX74, G155fsX197, S855X), Mean with SD, Two way ANOVA, Sidak’s post-hoc, \*\*\*\**P<0.0001.* **D**. Graphs showing that a very high percentage of iPS derived nociceptors (differentiation day 12: E55fsX74, G155fsX197, S855X) display CGRP and βIII-tubulin co-expression. Almost the totality of differentiated neurons (βIII-tubulin+) are positive for the nociceptor specific marker CGRP. No difference was observed between mutant and isogenic control cells or between WT and *Cdkl5^-/y^* murine DIV3 DRG cultures, where the percentage of βIII-tubulin and CGRP co-expressing neurons is about 40% of the total number. N: 3 independent experiments, Mean with SD, Multiple *t*-test, Two stage step-up method, *P>0.05*. **E**. Graphs showing the percentage of capsaicin responding cells compared to the total number of KCl responding DIV-42 iPS derived nociceptors (E55fsX74, G155fsX197, S855X) and WT murine DIV3 DRG neurons. Almost the totality of the iPS derived neurons respond to capsaicin equally between mutant neurons and their isogenic controls. Around 60-70% of DIV3 DRG neurons respond to capsaicin equally between WT and *Cdkl5^-/y^* cultures. N: 3 independent experiments, Mean with SD, Multiple *t*-test, Two stage step-up method, *P>0.05*.

**Supplementary Figure 3.**
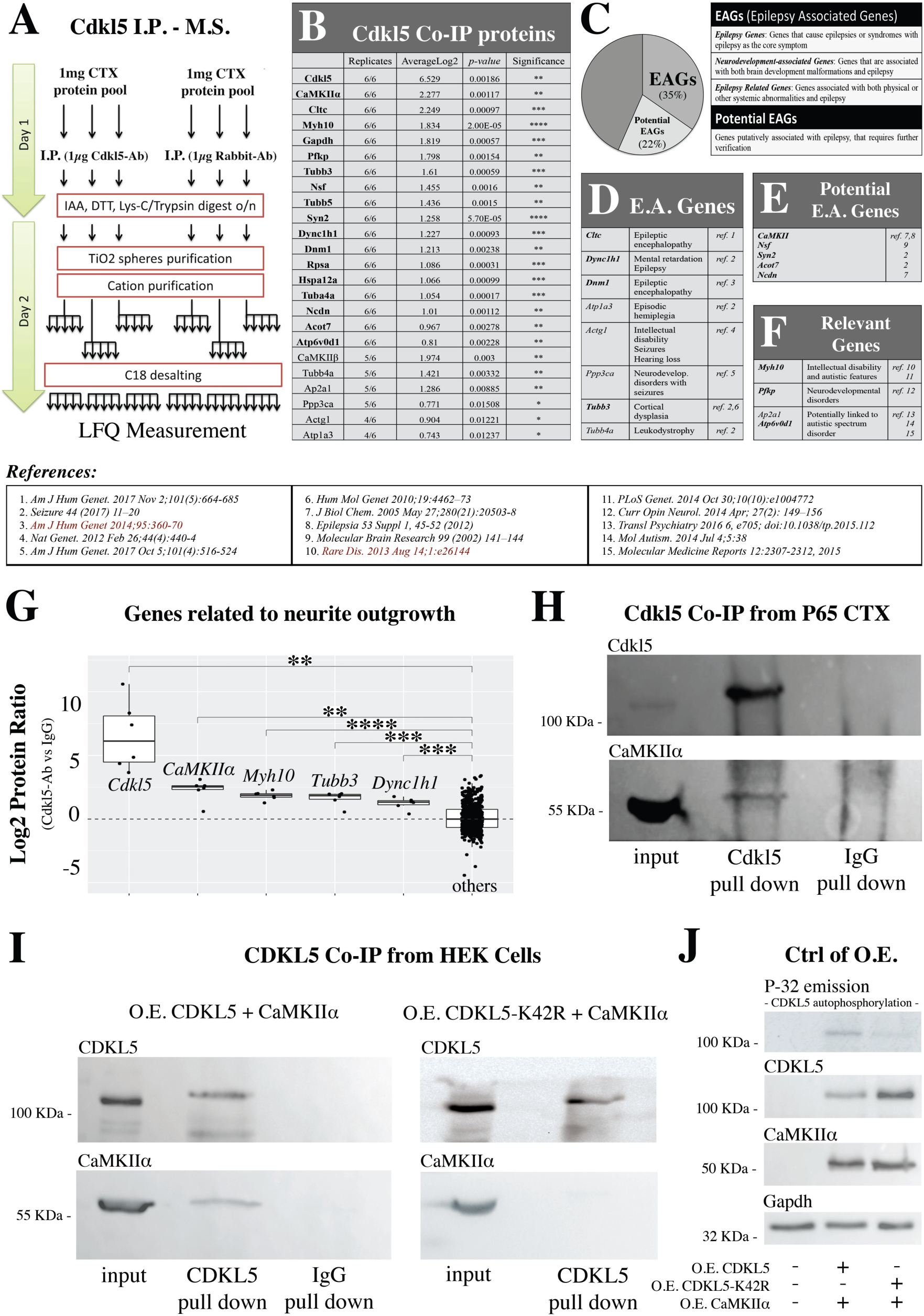
**A**. Schematic workflow illustrating the sample preparation and protein measurement by mass spectrometry after immunoprecipitation for Cdkl5 from the murine cortex. The peptides were labelled LIGHT (DimethLys-0 and DimethNter0) and HEAVY (DimethLys-4 and DimethNter4; IgG control). **B**. List of the top 23 proteins enriched in the Cdkl5-IP eluate. The statistical significance has been calculated comparing the Log2 ratio values from the Cdkl5 and the IgG eluate. N: 6 biological replicates, Parametric Two-tailed test, \*\*\*\**P<0.0001* \*\*\**P<0.001* \*\**P<0.01* \**P<0.05*. **C-F**. Classification of the Cdkl5-Co-IP proteins listed in B based upon their demonstrated or putative role in epilepsy. 35% of the Cdkl5-Co-IP proteins are “Epilepsy Associated Genes” (EAGs) and the 22% are potential EAGs (**D, E**). Some of the proteins are also related with intellectual disabilities, autistic features and other neurodevelopmental disorders (**F**). **G**. Log2 protein ratio and statistical analysis of the top ranked shows Cdkl5 putative interactors related to neurite outgrowth including CaMKIIα. N: 6 biological replicates, Parametric Two-tailed test, \*\*\*\**P<0.0001* \*\*\**P<0.001* \*\**P<0.01* \**P<0.05.* **H**. Western blotting showing co-immunoprecipitation of CaMKIIα after Cdkl5-pull down from adult murine cortical protein extracts. **I-J**. Representative western blotting showing co-immunoprecipitation of CaMKIIα after CDKL5-pull down from HEK-293 Cells overexpressing CaMKIIα together with CDKL5. The co-immunoprecipitation is impaired when CaMKIIα is co-transfected with the inactive form of CDKL5 (Kinase-Dead CDKL5-K42R) (**I**). The overexpression of the kinases is confirmed by Western Blot analysis, while the activity (autophosphorylation) of CDKL5 and CDKL5-K42R have been tested in vitro (Kinase Assay followed by SDS-PAGE) (**J**).

**Supplementary Figure 4.**
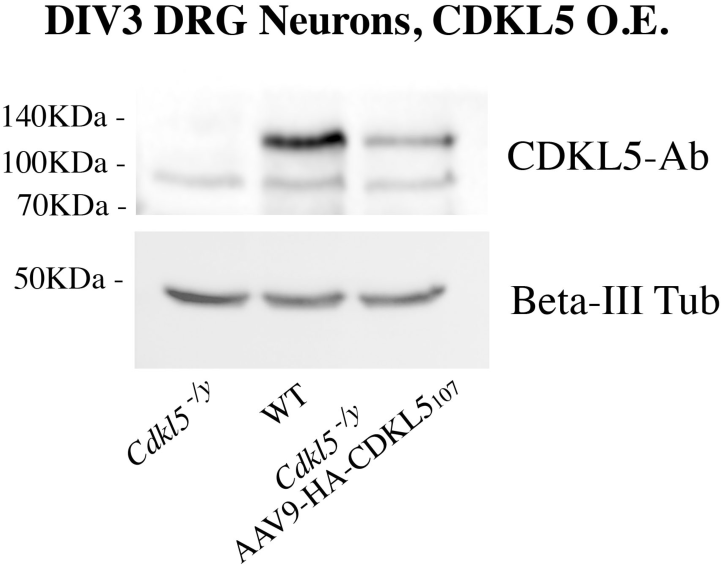
Western blotting showing the expression of CDKL5 in DRG cultures (DIV3) from WT and *Cdkl5^-/y^* mice, after AAV9-mediated overexpression of CDKL5. The neurons were infected (AAV9-ss-pTR-CBh-HA-CDKL5_107_) 24 hours after plating. βIII-tubulin was used as loading control.

**Supplementary File 1.** Shown is a list of proteins detected by mass spectrometry following immunoprecipitation for Cdkl5. Log2 normalized forward and reverse ratios as well as *p* values are provided.

